# Computational Characterization of Decision Making During Trans-saccadic Visual Perception

**DOI:** 10.1101/2025.10.07.680872

**Authors:** Pierre G. Gianferrara, Weiwei Zhou, Tyler A. Lesh, Timothy D. Hanks, Cyndi L. Edelman, Cameron S. Carter, Wilsaan M. Joiner

**Author notes:** Correspondence should be addressed to PGG.

## Abstract

When viewing the environment, humans execute fast sequential saccadic eye movements that disrupt the flow of visual information, yet still preserve stability in their visual percept. At the time of the saccade subjects often experience saccadic suppression of displacement—a failure to notice changes in the location of visual objects slightly shifted during the eye movement. Although well studied, the weighing of sensory information and the extent to which this integration influences visual perceptual judgments is still poorly characterized and understood. Here we used a drift diffusion modeling (DDM) framework to systematically examine the extent to which sensory information biased perceptual judgments when detecting trans-saccadic shifts of visual targets. Healthy human participants (*N* = 29, 20 female) completed a visual perception task in which a visual target was shifted following a cued saccadic eye movement (4° or 8°). Trans-saccadic target displacements occurred up to ±2.5° along the horizontal axis, and participants reported the shift direction with a button press response. We present a version of the DDM that takes the visual error as input to simulate offsets in evidence accumulation during decision-making. The model provides an explanatory framework for the integration of sensory information, capturing the temporal dynamics of evidence accumulation that explain the range of perceptual biasing and response timing effects observed across participants.

**SIGNIFICANCE STATEMENT:** Maintaining a stable visual percept between saccadic eye movements relies on the integration of externally-generated sensory signals and internally-generated signals based on the executed movements. In this work, we provide a computational framework (a modified drift diffusion model) to quantitatively establish the relative contribution of visual error signals to perceptual judgments and decision-making behavior following saccades. We show that greater reliance on visual error leads to increased biasing effects on response accuracy and manual reaction time. These effects are characterized in the DDM by a visual error-based modulation of the drift rate, representing evidence accumulation. Collectively, the results provide novel measures that capture the influence of sensory information on decision-making and explain the inter-subject differences observed in trans-saccadic perception.

## INTRODUCTION

One remarkable property of the visual system is its ability to rapidly sample inputs from the surrounding environment and construct a seemingly coherent and stable perceptual representation of the world (Wurtz, 2008; Higgins and Rayner, 2015). Visual sampling occurs through fast-paced sequential eye movements (e.g., several saccades/second), which interrupt the continuous stream of incoming visual input (Wurtz et al., 2011). To correct for apparent perceptual motion due to retinal shifts, the visual system leverages compensatory mechanisms which integrate information across *externally* and *internally* generated signals (Feinberg, 1978; Hauser et al., 2011; Frith, 2015; Ondobaka et al., 2015). In the brain, externally generated, largely sensory, signals must be distinguished from internally-generated signals, which are theorized to arise from internal copies of issued motor commands (i.e., corollary discharge; Joiner et al., 2010; Kawato, 1999; Kawato et al., 2021; Wolpert et al., 1998; Wurtz, 2008).

During saccadic eye-movements, it has been observed that subjects often fail to notice changes in the location of objects, particularly with small changes in amplitude. This phenomenon (saccadic suppression of displacement, SSD) has been attributed to the brain’s decreased perceptual sensitivity around the time of the saccade (Takano et al., 2020). In addition, seminal findings by Bridgeman et al. (1975) and Westheimer (1984) found that retinal acuity tended to decrease as a function of the saccade amplitude. Based on Bayesian models of trans-saccadic integration, these results have been recontextualized as optimal inference mechanisms whereby the brain may be maintaining a sense of spatial constancy by pooling information from retinal locations, velocity signals, and a proprioceptive sense of the eye position (Gysen et al., 2002; Niemeier et al., 2003; Atsma et al., 2016).

Examining the mechanisms underlying the integration of foveal and extraretinal signals requires understanding how perception is impacted by the saccade magnitude and extent SSD might be affected by the visual objects’ retinal shifts. The displacement detection task from Bridgeman et al (1975) provides an excellent paradigm through which to study this perceptual process. In this task, participants execute saccadic eye movements towards an initial peripheral target. While participants make the saccade, the target is extinguished and reappears at a slightly shifted location. Subjects then report the perceived direction of the target shift (Bridgeman et al., 1975; Deubel et al., 1996, 1998; Bansal et al., 2015).

Although the psychophysical effects of SSD have been referenced in prior work (Deubel et al., 1996, 1998; Niemeier et al., 2003; Atsma et al., 2016; Takano et al., 2020), it is currently unknown to what extent the decision making process is altered by the processing of externally-generated signals. Here, we leveraged a computational framework, the drift diffusion model (DDM; Ratcliff, 1978), to gain insight into the decision-making process underlying the perceptual report (Ratcliff and Smith, 2004; Ratcliff and McKoon, 2008; Mulder et al., 2014). One advantage of the DDM is that it can both model choice and reaction time data (Mulder et al., 2014; Cerracchio et al., 2023; Forstmann and Turner, 2024).

Within the displacement detection task, DDM simulations may help determine how participants’ weighing of sensory information throughout evidence accumulation may result in a behavioral bias during perception (Witthoft et al., 2018; Cerracchio et al., 2023). We focused on the DDM’s account of the visual error (VE) contribution to decision-making, where VE was defined in terms of the visuospatial offset between the eye endpoint location and the visual target location. In cases of perceptual ambiguity, we hypothesized that a greater reliance on the VE during decision-making would lead to greater biases in perceptual judgments and reaction time profiles whereas a lower reliance on the VE would result in more accurate perceptual judgments due to the integration of extraretinal signals (Lange et al., 2021; Goris et al., 2024).

We tested this by comparing a modified version of the DDM (with a bias term that offset the drift rate based on the VE) to a baseline DDM with manual choice and reaction time symmetry. We show how inputting VE coefficients into the DDM provides a more granular representation of individual evidence accumulation speed biases across target shift directions and saccadic amplitudes. In terms of inter-individual differences, we demonstrate how perceptual biases that are more congruent with VE-based predicted responses tend to be linked to systematic behavioral patterns of asymmetry in psychometric and RT profiles. Overall, our modified DDM-based framework provides a novel method to computationally capture the influence of sensory information on decision-making, and explain perceptual biasing and response timing effects.

## MATERIALS AND METHODS

### Participants

Thirty healthy participants took part in this study. One participant was excluded due to poor eye-tracking (i.e., no variation in the VE across target shifts). Subsequently, we report data from 29 participants aged 18-41 years-old (*M* = 22.62 years-old; *SD* = 5.39 years). Twenty participants were female, and 9 participants were male. All had normal or corrected vision. Informed consent was obtained from every participant, and study protocols were approved by the Institutional Review Board at the University of California, Davis. All study procedures were conducted at UC Davis Imaging Research Center in Sacramento, CA.

### Apparatus

Participants were seated in a darkened room with their head stabilized by a chin and forehead rest, 100 cm in front of a 1980 x 1200 Dell U2415 monitor with 60 Hz refresh rate (Dell Technologies Inc., Round Rock, TX). Manual choices were registered on a button box at the desk. Fixations and saccades of the right eye were recorded with the Eyelink II eye tracker (head-mounted binocular eye tracker, 1000 Hz temporal resolution, 0.01° spatial resolution, Eyelink CL version 5.15 source, Eyelink GL camera version 1.2; SR Research Ltd., Mississauga, ON, Canada) and E-Prime 2 software (Psychology Software Tools Inc., Sharpsburg, PA). Stimulus presentation and manual button box response data were acquired in E-Prime 2. Stimuli presented were either a black fixation cross or a filled black circular target on a gray background. At the start of the experiment, a five-point calibration and validation were performed.

### Task and Procedure

The task was adapted from a previous trans-saccadic visual perception experiment (Bansal et al., 2018). Each trial started with a fixation cross (0.3° in extent & 8.5 mm height and width) presented at the center of the screen. Participants fixated the cross for a random duration between 1200 and 1800 ms. A circular target (8.5 mm diameter) appeared in either the left or right direction from fixation and required a saccade of amplitude 4° or 8°. As soon as the eye-tracker determined that participants’ eye position was no longer in a 3.2° square window centered around the fixation cross, the target disappeared (i.e., blank screen) for a time period of 250 ms, and subsequently reappeared at one of 11 shifted locations ranging between - 2.50° and + 2.50° (see Fig. 1A for illustration of the task).

**Figure 1.**
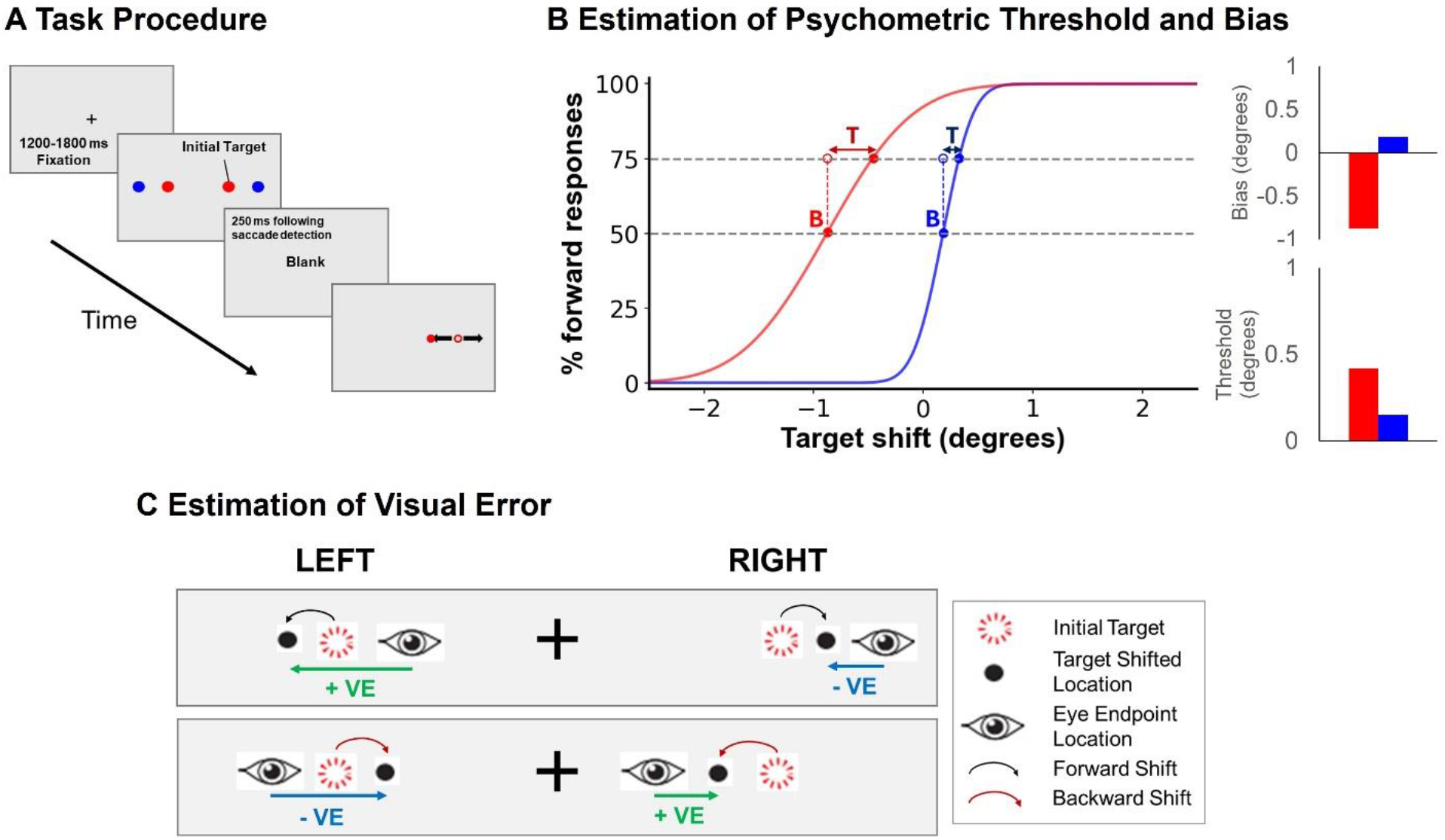
Description of the Trans-saccadic Shift Detection Task. (A) Overview of the task procedure. Trials started with a fixation cross. An initial target then appeared on either the left or right side of the screen and required a 4° or an 8° saccadic eye movement. Following a 250 ms blank screen, the target reappeared at a shifted position ranging from 2.5° to the left to 2.5° to the right. (B) Overview of the psychometric threshold and bias. The proportion of forward button press responses is plotted against the target shift magnitude. Button press responses were fitted with a beta binomial model to determine the psychometric function which were the basis of the psychometric bias and psychometric threshold. The psychometric bias (indicated with a ‘B’) corresponds to the point on the curve at which there were 50% forward responses. The psychometric threshold (indicated with a ‘T’), on the other hand, corresponds to the step, in degrees, that is required to go from 50 % forward responses to 75% forward responses. On the plot, the red curve corresponds to an example psychometric function with a negative bias and a high threshold. The blue curve, on the other hand, corresponds to an example psychometric function with a slightly positive bias and a low threshold. (C) Visual error (VE) description. A target shift could take place in the forward direction (i.e., away from the fixation cross; black curved arrow) or it could take place in the backward direction (i.e., back towards the fixation cross; red curved arrow). The VE indicates the difference between the eye position and the shifted target position. Consistent with the target shift direction, a positive (forward) VE corresponds to a case in which the shifted target lies more forward relative to the eye position. A negative (backward) VE corresponds to a case in which the shifted target lies more backward relative to the eye position.

Non-zero target shifts were uniformly distributed, but there were twice as many trials that did not involve any visual displacement of the initial target (i.e., zero shift) (see target shift axis on Fig. 2 and 4). In the experiment, we compared forward vs. backward target shifts. Forward shifts refer to visual targets that reappeared further away from the fixation cross. Backward shifts refer to the visual targets that reappeared closer to the fixation cross (see Fig. 1C for illustration of forward (top panel) and backward shifts (bottom panel)). Across all blocks, there were 80 instances of zero shifts, and 40 instances of each of the 10 non-zero target shifts. For each target shift, half of the trials involved initial targets appearing to the left or right of the fixation cross, and half of the trials required 4° or 8° saccades. Participants were instructed to make a manual decision (pressing the ‘left’ or ‘right’ labeled buttons) to indicate the target shift direction as quickly as possible, without sacrificing accuracy. Forward responses refer to button presses whose indicated direction (‘left’/‘right’) was *congruent* with the initial saccade direction (‘left’/‘right’) and are represented with a positive angle (in °). Conversely, backward responses refer to button presses whose indicated direction was *incongruent* with the initial saccade direction and are represented with a negative angle (in °).

**Figure 2.**
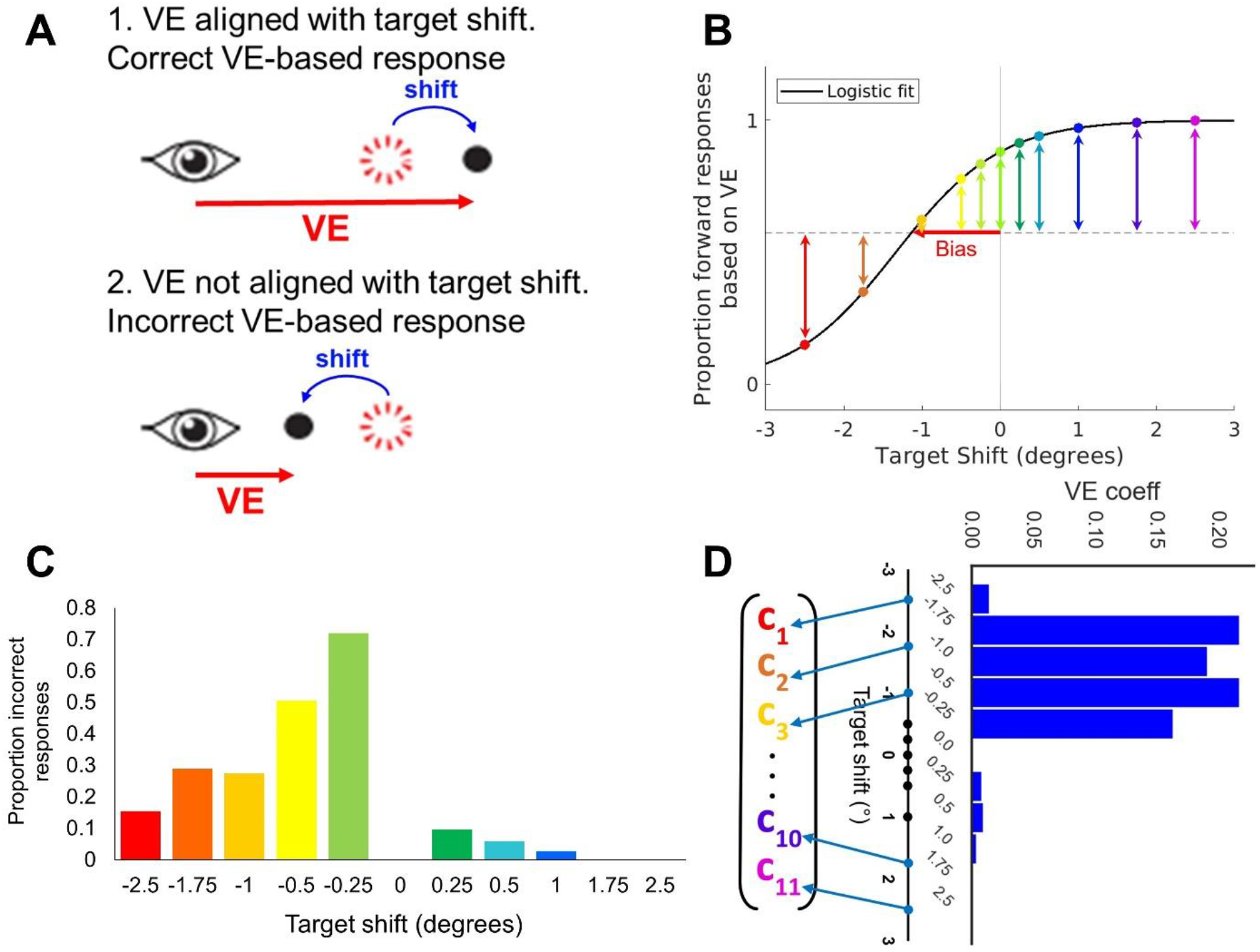
Extraction of VE-related coefficients indicating perceptual biasing effect on decision-making across target shifts. (A) The diagrams provide a visual illustration of a correct VE-based response with congruency between the target shift and VE sign (1) and an incorrect VE-based response with incongruency between the target shift and VE sign (2). (B) Proportions of *predicted* button press responses that would be in the forward direction solely based on the experienced VE for an example subject. The black line is the logistic curve fit to the respective proportions. The colored vertical arrows indicate the distance between each point for each target shift and the mid-point between the logistic function’s upper and lower asymptote. This indicates the inverse likelihood that a participant may be incorrectly influenced by the VE at that particular target shift. The general VE bias consists of the shift of the logistic curve along the horizontal axis relative to 0 degrees (indicated with a vertical gray line) at the point where the logistic curve crosses the dashed line indicating 50 % (0.5) forward responses and is shown in red. (C) Proportion of incorrect button press responses across target shifts for the same example subject. (D) VE coefficients integrating information from the proportions of incorrect responses and VE-predicted forward responses across target shifts. A high VE coefficient indicates a high propensity for decision-making bias (e.g., a target shift of -0.5°) whereas a low VE coefficient indicates a low propensity for bias (e.g., a target shift of +1.0 °).

Prior to the start of the experiment, participants were given three attempts at a 10-trial practice run to familiarize themselves with the task. Target shift magnitudes from the practice run had the same frequency distribution as the main experimental trials. Overall, feedback on task accuracy was provided at the end of each practice run, but no feedback was provided after trials from any of the 10 blocks. There were 480 trials throughout the experiment (48 trials separated into 10 blocks). The task lasted 30 minutes on average.

### Experimental Design and Statistical Analysis

We report below all experimental measures that were extracted, analyses and computational models that were run, and statistical tests that were conducted. For each statistical test, we considered the confidence intervals around our main estimates to provide an indication of uncertainty. All such confidence intervals are reported in the Statistical Table (see Table 1).

**Table 1.**
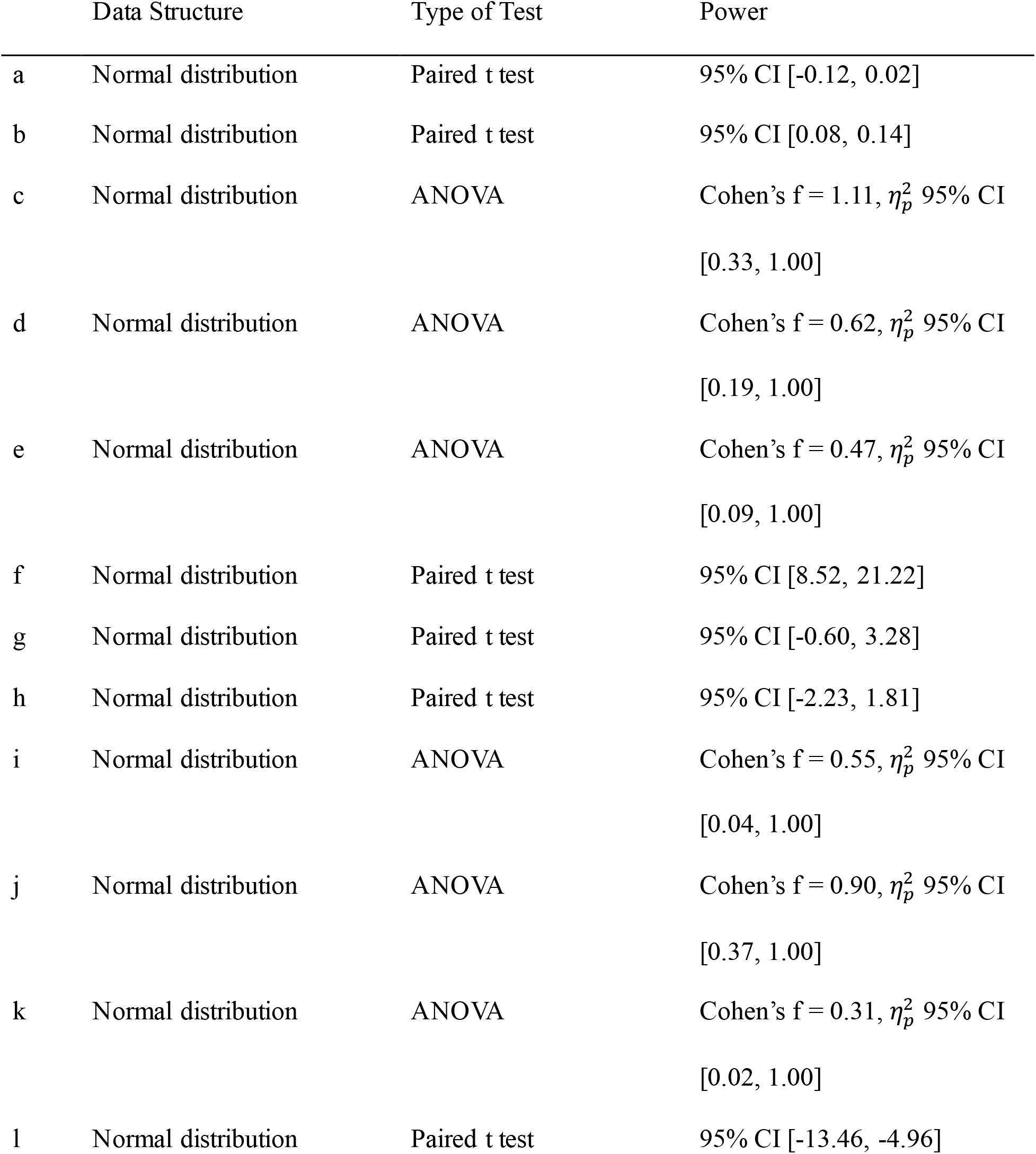
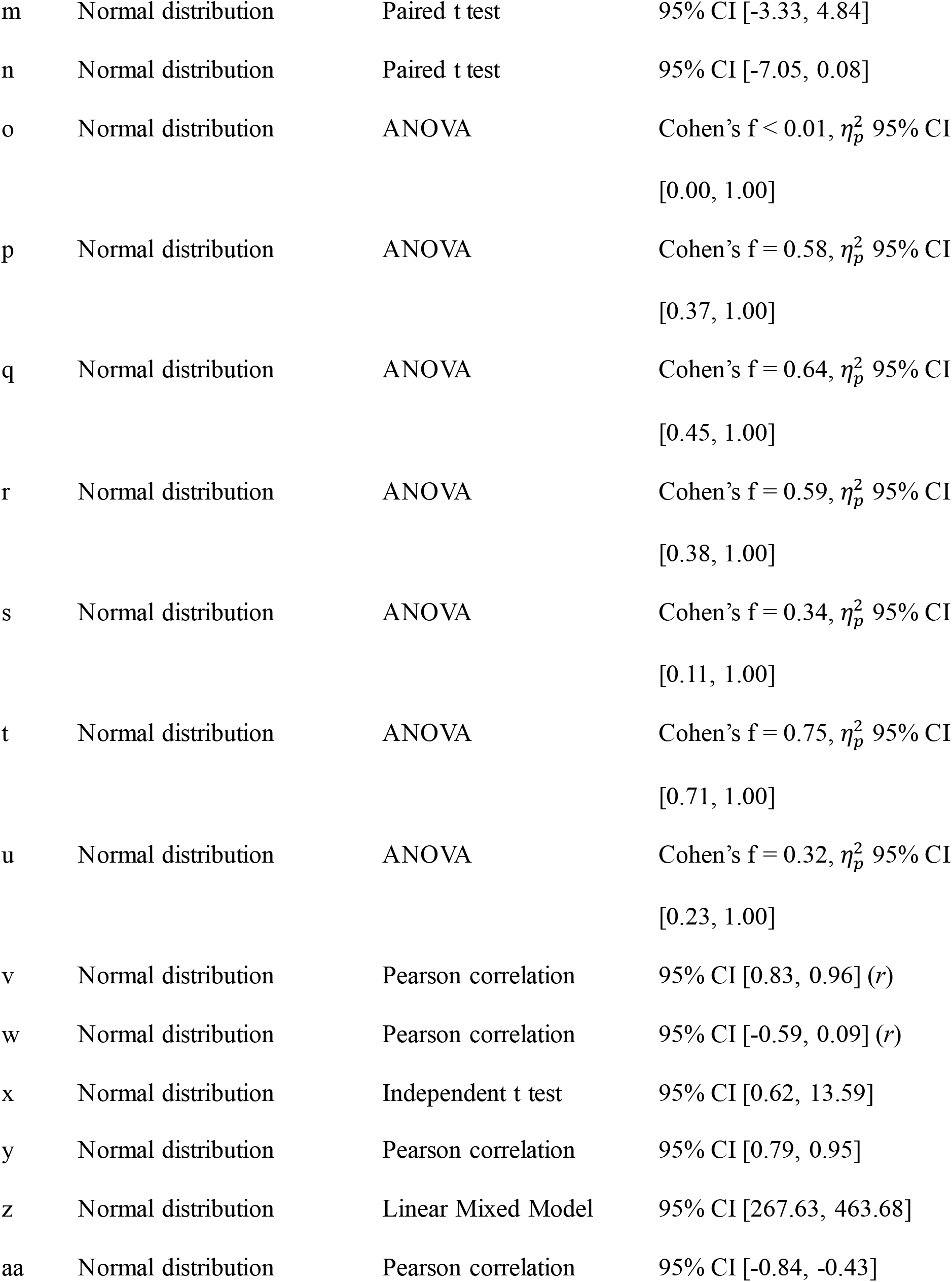
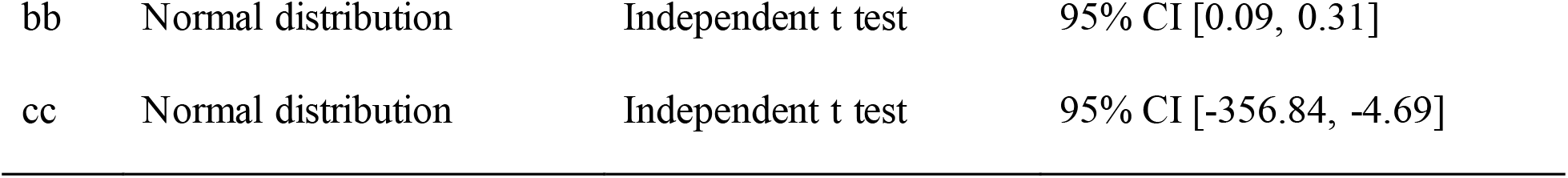
Statistical Table.

#### Behavioral measures

Accurate responses consisted of button presses whose indicated direction (‘left’ and ‘right’) matched the target shift direction (‘left’ vs. ‘right’), and were coded with a 1 whereas inaccurate responses were coded with a 0. In the case of the 0° target shift, either button press was considered correct (and coded with a 1) since no target shift occurred. Choice data were computed in terms of % forward responses (based on participants’ button presses) relative to the total number of trials for that target shift direction and amplitude. We reported the group mean accuracy (%) and standard error of the means across all target shifts for trials involving a saccadic eye movement of 4° and 8° amplitudes separately.

Reaction time (RT) was the time interval between when the target reappeared, and the button press onset. We first computed the mean manual RT across trials with zero target shift and reported the RT at all other target shifts in terms of a percentage relative to the zero target shift RT. Zero target shifts represent the case in which participants had to guess because the target did not move. Normalization of RT by the zero target shift was carried out within subjects. We reported the group mean manual RT (%) and standard error of the means across all target shifts for trials involving a saccadic eye movement of 4° and 8° amplitude separately.

#### Psychometric analyses

In order to capture participants’ response profiles across target shifts, we characterized participants’ perceptual sensitivity to target shifts with psychometric curves and a posterior distribution estimated with Bayesian methods and a beta-binomial model fitted to manual choice (proportion of forward responses; Schütt et al., 2016). In terms of the psychometric curve itself, psychometric sensitivity was quantified in terms of the percentage of forward responses as a function of the target shift (see Fig. 1B). Two measures were extracted from individual participants’ psychometric curves—the *psychometric bias* and the *psychometric threshold* (see Fig. 1B for illustration of both).

The psychometric bias (labeled with a ‘B’ on Fig. 1B) corresponds to the point at which the participant reported that the target jumped forward 50% of the time (i.e., at chance level). A positive psychometric bias indicates that the participant tended to report backward responses for small forward target shifts. The blue curve in Fig. 1B illustrates an example of a slight positive psychometric bias. Conversely, a negative psychometric bias indicates that participants tended to report forward responses for small backward target shifts. The red curve in Fig. 1B illustrates an example of a negative psychometric bias.

The psychometric threshold (labeled with a ‘T’ on Fig. 1B) corresponds to the target shift difference (or step) required to transition from 50% forward responses to 75% forward responses. This measure corresponds to a participant’s difficulty in perceiving trans-saccadic shifts. The psychometric threshold tends to be highest when participants make numerous erroneous responses at the transition point between backward and forward target shifts, thus resulting in a flatter psychometric curve and indicating low perceptual sensitivity. This corresponds to the red psychometric curve in Fig. 1B. Conversely, the psychometric threshold tends to be lowest when participants make very few erroneous responses, thus resulting in a steeper psychometric curve and indicating high perceptual sensitivity. This corresponds to the blue psychometric curve on Fig. 1B. In MATLAB, psychometric functions were fit with the *psignifit* function (Schütt et al., 2016). See Table 2 for description of key psychometric measures.

**Table 2.** List of Key Experimental Measures.

| Variable | Definition | Purpose |
| --- | --- | --- |
| Psychometric bias | Point at which participants reported forward target jump 50% of the time | Quantify shift of psychometric function along horizontal dimension |
| Psychometric threshold | Target shift difference (i.e., step) required to transition from 50% to 75% forward responses | Inversely quantify degree of perceptual sensitivity in the trans-saccadic task |
| VE-based forward response | Prediction of forward button presses that would result from VE | Establish psychometric profile solely based on VE information |
| VE bias | Estimation of the inflection point based on VE logistic curve (Eq. 2) | Quantify shift of the VE logistic profile |
| VE coefficient | Proportion of incorrect perceptual responses divided by distance from VE-predicted forward response mid-point | Quantify propensity of VE-based perceptual inaccuracies |
| DDM bias | Shift of the starting point in the DDM | Simulate asymmetry in reaction time and psychometric profile |
| VE coefficient centroid | Center of gravity of the distribution of VE coefficients | Extent of biasing effect of VE on decision-making |

#### Saccadic and visual error measures

Perception was only assessed in the horizontal plane as previous research has shown that there are no significant trans-saccadic perceptual differences between horizontal and vertical movements (Bansal et al., 2015). Successful trials required a detectable initial saccade (eye velocity ≥ 50°/s and acceleration ≥ 8000°/s²) and a response between 200 ms and 1,200 ms of the shifted target’s appearance (Joiner and Shelhamer, 2006; Joiner et al., 2007). Saccade initiation was detected when the eye position exited the fixation window. Included saccades were all initiated within the fixation window, had a saccade timing onset within 300 ms after the appearance of the first target, with valid eye fixations at the time of the target reappearance.

We computed the gain to quantify the amount subjects undershot or overshot the initial target location. Saccade gain was calculated relative to the target in terms of the location of the initial saccade and the initial target location:

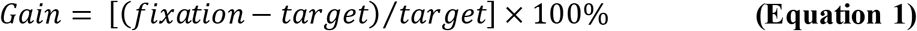

Overshoots were defined as saccades whose gain exceeded 100% and undershoots were defined as saccades whose gain was lower than 100%.

Visual error (VE) corresponds to the difference between the eye endpoint location and the shifted target location along the horizontal axis. The VE’s sign indicates the position of the eye fixation relative to the shifted target. A positive VE occurred when the target’s shifted location was in the forward direction and a negative VE occurred when the target’s shifted location was in the backward direction.

VE information was subsequently used to make predictions of the manual responses that would directly result from the experienced VE. Such information was integrated across target shifts for each of the two saccade amplitudes (4° and 8°) and used to determine the general VE bias. Estimation of this bias relied on a logistic function fit to VE data, which was defined as the point corresponding to 50% VE-predicted forward responses (i.e., 0.5 on the logistic function; see Fig. 1B) and refers to the shift of the logistic function along the backward/forward direction. The logistic function was defined as:

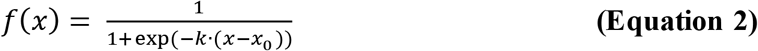

where *k* refers to the growth rate (higher *k* indicates a faster switch), *x*_0_ refers to the inflection point, 2 is the range, and -1 is the fixed intercept to ensure the function ranges from -1 and 1 as its two fixed asymptotes. We were particularly interested in estimating *x*_0_ as a measure of VE bias, which corresponds to the point at which there is 50% VE-predicted forward responses.

In order to obtain insight into the specific target shifts at which there was a discrepancy between the shift’s direction and the VE, we computed the VE-shift. Specifically, if a subject makes a saccade directly to the initial target, the expectation was that a visual target being shifted in the forward direction would result in a forward VE relative to the eye endpoint location (see Fig. 2A). Conversely, a visual target being shifted in the backward direction would be expected to result in a backward VE relative to the eye endpoint location. For each target shift, we calculated the % VE-shift congruency in terms of the proportion of trials that resulted in the expected sign congruency relative to all trials at that specific target shift. We did this separately across saccade amplitudes of 4° and 8°. See Table 2 for description of key experimental measures.

#### VE coefficient estimation & centroid

We quantified the degree to which an individual’s perceptual judgments relied upon VE by computing the distribution of VE coefficients based on the frequency of perceptual decision-making errors and the probability of such errors based on the VE (see Fig. 2D). For the latter, we determined the probability of a forward response on each trial based solely on the VE (Fig. 2B). Next, for each target shift, the corresponding VE coefficient was computed in terms of the proportion of incorrect responses (see Fig. 2C) divided by the distance between the VE-based predicted forward response on the logistic function (see Fig. 2B) and the mid-point of the VE-predicted forward response logistic function upper and lower asymptote (see Fig. 2B). Note that only one distribution of VE coefficients was estimated based on the average VE-predicted forward responses across all trials (both 4° and 8°; Fig. 2B). Overall, participants were more likely to make VE-driven perceptual judgment errors when the sign of the VE vector was inconsistent (i.e., not aligned) with the direction of the target shift (see Fig. 2A for illustration).

We leveraged the distribution of VE coefficients to estimate the extent of the biasing effect of the VE on decision making. This measure consisted of the center of gravity, or centroid, of the VE coefficient distribution:

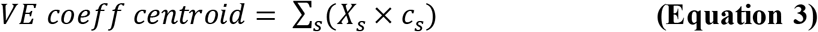

where *X_s_* indicates a signed target shift in the backward or forward direction, and *c_s_* is the VE coefficient corresponding to the target shift *X_s_*. To assess the statistical robustness of the VE coeff centroid, we computed the signal-to-noise ratio (SNR):

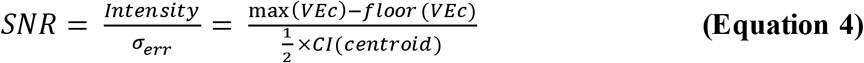

where *max(VEc)* is the peak height of the VE coefficient distribution, *floor(VEc)* is the median of the lower half of the ranked VE coefficients, and *CI*(*centroid*) is the confidence interval estimated after bootstrapping 2,000 times by resampling with replacement pairs of VE-predicted forward responses and response accuracy across all experimental trials for each target shift separately.

#### General description of the drift diffusion model (DDM)

The objective of the DDM was to simulate the evidence accumulation process leading to perceptual judgments, which corresponds to a noisy process (see Extended Data Figure 3-1 for illustration of the DDM). DDM model inputs consisted of an 11 x 8 matrix, which included the target shifts, the forward and backward manual reaction time means and standard error of the means, the number of forward responses, and the total number of trials corresponding to a given target shift. An additional input captured VE-related biasing effects for each target shift. This corresponded to the distribution of VE coefficients (see Fig. 2D). See Fig. Extended Data Figure 3-2 for an illustration for the DDM input matrix.

The model progressively accumulates momentary evidence towards an upper (“A”) or lower bound (“B”; see Extended Data Figure 3-1). Each bound indicates one out of two possible choice options (i.e., “left” vs. “right” target shift report). Positive evidence leads to a greater probability of selecting choice “A”, whereas negative evidence leads to a greater probability of selecting choice “B”. Mathematically, the probability of exceeding the decision threshold A before B is represented by the following equation:

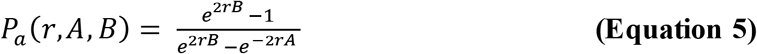

The mean time (*T*) to bound A is represented by:

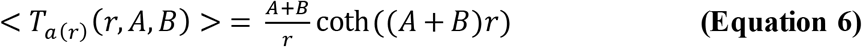

There were four main parameters that were estimated by the DDM with the negative log likelihood estimation technique: drift rate coefficient (*r*), decision bound (*DB*), non-decision time (*NDT*), and starting bias (B; Palmer et al., 2005; Ratcliff and McKoon, 2008; Smucny et al., 2023a). At each step along the decision-making process, new evidence is accumulated and drawn from a unit-variance Gaussian distribution with mean set by the drift rate, which we designate *r* in the equations above and corresponds to a linear function of the stimulus input. The *DB* indicates the choice that is selected (“A” vs. “B”), and the corresponding decision time is determined by how long it took the DDM to pass the decision threshold during evidence accumulation. Therefore, the *NDT* parameter estimate indicates a proportion of the manual reaction time that is not directly related to the evidence accumulation process. Finally, the starting bias *B* represents an initial offset that favored one of the two possible decisions by shifting the *DB* in the forward or backward direction. The bias was specifically quantified by the extent to which the starting point (0 point on Extended Data Figure 3-1 & Fig. 3A) was shifted upwards towards the “A” decision bound (i.e., positive bias), or downwards towards the “B” decision bound (i.e., negative bias; see Fig. 3A for an illustration of a downwards bias). For the purposes of these computational simulations, the DDM was applied within-subjects across all target shifts for all trials corresponding to a given saccadic eye movement amplitude (4° vs. 8° trials). All parameters were fit by minimizing the negative log likelihood of the data (i.e., maximizing likelihood of data; Smucny, Hanks, Lesh, & Carter, 2023).

**Figure 3.**
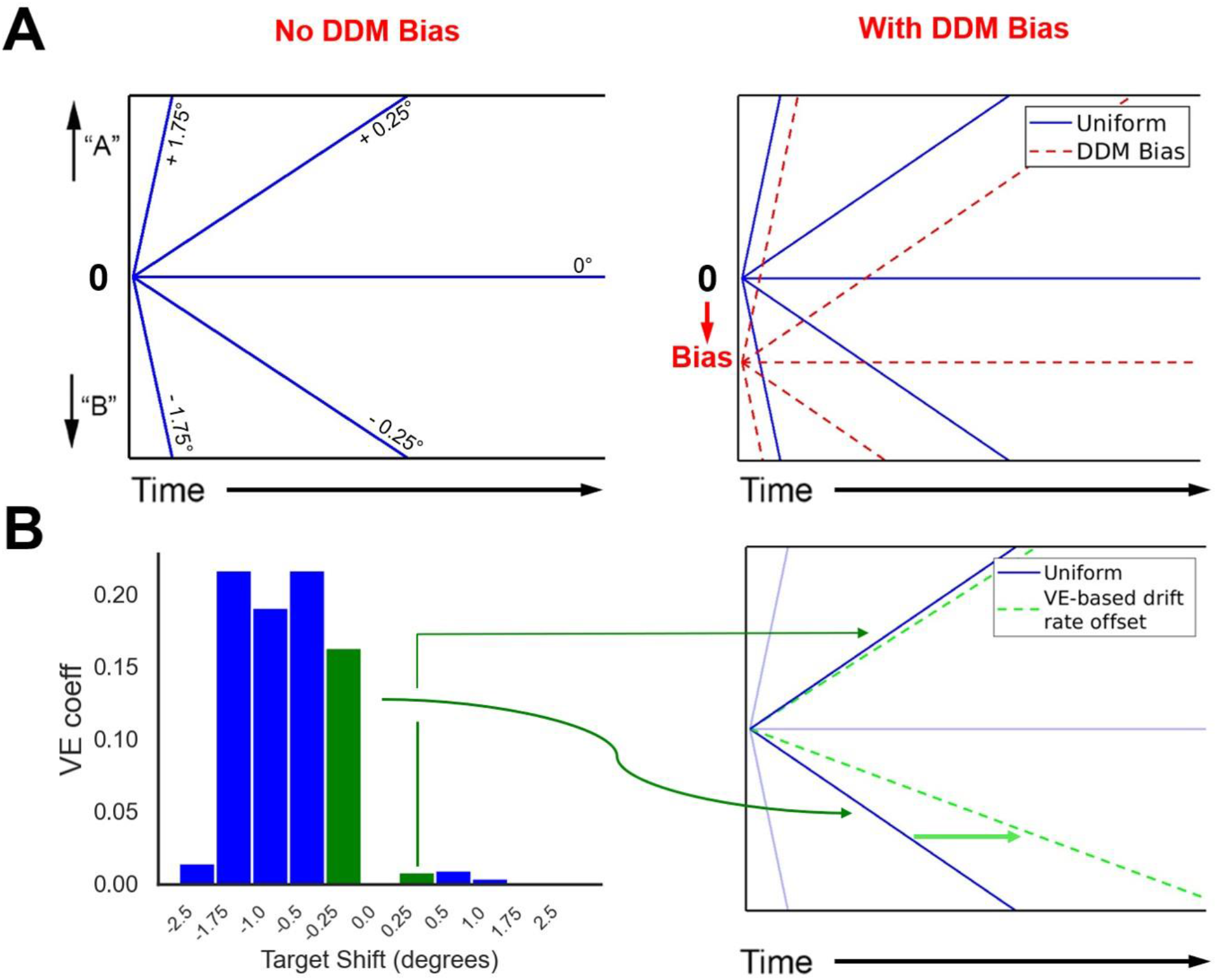
Visual Representation of the DDM Models and biases. (A) Illustration of the DDM starting bias. On the left, the uniform model assumed a symmetric architecture across target shifts in the forward and backward direction. The model accumulates momentary evidence in either direction (“A” vs. “B”) until a decision bound is reached. On the right, the model with a DDM bias involves a shift of the starting point (indicated with a 0), which creates reaction time asymmetry across the drift rates corresponding to the forward vs. backward direction. (B) Illustration of the VE-related biasing effect on the drift rate in the model that includes a DDM bias. Drift rates are additionally offset at the level of the inputs proportionally to the VE coefficients across target shifts. Across target shifts, a higher VE coefficient corresponds to a higher drift rate offset.

To simulate potential biasing effects due to the VE, we modulated the DDM *input_s_* across target shifts denoted with an *s*, which were expressed as a function of the VE coefficients across target shifts:

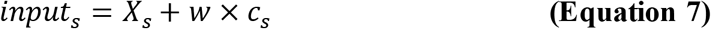

where *X_s_* indicates a target shift, *c_s_* denotes the corresponding VE coefficient, and *w* corresponds to an additional DDM VE-related parameter that is determined with the negative log likelihood estimation method to obtain the best fit with data. Since each target shift’s drift rate is directly proportional to the DDM inputs and thus the target shift’s corresponding VE coefficient, this resulted in an offset of the drift rate as illustrated on Fig. 3B. Note that there were unique *w’*s (i.e., DDM VE weight) for 4° and 8° saccade amplitude trials.

We compared our main DDM to a reference baseline model that we refer to as the “uniform DDM.” This uniform DDM differed from the modified VE-driven DDM described above in two ways. First, the uniform model did not include any starting bias meaning that the forward and backward target shifts with the same magnitude required the same time for participants to reach the forward decision threshold “A” and the backward decision threshold “B” (see Fig. 3A). Second, the uniform model did not modulate the input drift rate according to the VE coefficient distribution as indicated on equation 7. We did this to quantify the extent to which participants’ manual reaction time profiles differed from what would be expected from a symmetric DDM with no bias.

#### Statistical relationship between VE coefficients and RT

To test whether the VE coefficients could adequately capture the decision-making process unfolding over time, we estimated participants’ reaction time (RT) profiles based on the VE coefficients as part of a linear mixed effects model (LMEM). In the model, we set the VE coefficient as the main fixed predictor and added three random intercepts to account for variability across participants, saccade amplitudes, and target shifts. In R, the LMEM was defined according to the following equation: *Y_RT_* = *γ*_1_ *X_VEc_* + *Z*1*_subject_* + *Z*2*_amplitude_* + *Z*3*_s_*_ℎ*ift*_. We imported and utilized functions from the lme4 (Bates et al., 2015) and lmerTest packages (Kuznetsova et al., 2015) to fit the linear mixed-effects model. The model was written as lmer(RT ∼ VE_c + (1|subject) + (1|amplitude) + (1|target_shift)). The restricted maximum likelihood (REML) technique was applied to each model and we obtained a p-value for the *β* estimate. We also reported the 95% confidence interval around the *γ_1_* coefficients corresponding to VE *β* by performing the ‘confint’ *R* function with the Wald method.

#### Additional statistical analyses

All statistical analyses were performed in *R* version 4.1.3. For psychometric analyses, paired *t-*tests were performed on the estimated psychometric bias and threshold across saccades amplitudes 4° and 8°. We estimated correlational effects between the psychometric bias and VE logistic bias (i.e., from VE-predicted forward responses) with Pearson correlation (*r*) in the backward (negative psychometric bias) and forward (positive psychometric bias) separately. To test for potential differences in response accuracy, reaction time, and shift-VE congruency levels, we applied a 2×11 two-way analysis of variance (ANOVA) with repeated-measures and a Greenhouse Geisser correction. The factors were the saccade amplitude (4° and 8°) and target shift (11 possible shifts). For each ANOVA, and depending on significance levels, post-hoc pairwise *t-* tests were used with a Bonferroni correction. To test for VE bias differences across saccade amplitudes, we used a paired *t*-test on the group logistic fits’ inflection point *x_0_* across trials with 4° vs. 8° saccadic eye movements.

Differences in manual RT profiles across the uniform DDM and VE-driven DDM were evaluated with a χ^2^ statistic such that RTs from the uniform model consisted of the expected RTs and RTs from the VE-driven model consisted of the observed RTs. A low χ^2^ signified a symmetric RT profile and a high χ^2^ signified an asymmetric RT profile. Relationships between the VE coeff centroid (i.e., equation 3) and the psychometric bias and RT χ^2^ estimates were assessed across participants with Pearson correlation (*r*). Relationships between DDM parameters (drift rate *r*, *B*, and *NDT*) and experimental variables (psychometric threshold, VE coeff centroid, and RT) were also evaluated with Pearson correlation (*r*). Accuracy, VE coefficient centroid, and RT asymmetry differences across positive and negative psychometric groups were assessed with independent *t*-tests.

### Code Accessibility

All code, software, and data that were used and described in this paper are freely available online at https://doi.org/10.6084/m9.figshare.30090127 (Gianferrara & Joiner, 2026). All code is available as Extended Data 1. DDM scripts were written in MATLAB R2022a and are used to simulate the evidence accumulation process leading to perceptual judgments in the trans-saccadic task. The code and data have been deposited in Figshare under repository name “Computational Characterization of Decision Making During Trans-saccadic Visual Perception” and is licensed as CC BY 4.0.

## Results

The main objective was to use the DDM computational modeling framework to characterize the evidence accumulation process in the displacement detection task. We aimed to determine to what extent a greater contribution of the VE to perceptual judgments would result in a biasing effect on and greater asymmetry in perceptual responses and manual reaction time profiles. We did this across individuals by considering a uniform DDM and a DDM modified by the VE. Across all results and for each statistical test, Table 1 reports the corresponding confidence intervals as uncertainty estimates.

### Psychometric and Manual Choice Measures

Fig. 4 presents the behavioral results from the visual perception task. Psychometric curves were fitted to individuals’ proportions of forward responses across target shifts. No significant differences across 4° and 8° group mean biases were found, *t* (28) = - 1.47, *p* = 0.15^a^. In terms of the psychometric threshold, there was a significantly larger threshold for the 8° trials compared to the 4° trials, *t* (28) = 7.21, *p* < 0.001^b^. Consistent with previous work (Bansal et al., 2015; Bansal and Joiner, 2021), this result indicates lower sensitivity in the detection of target shifts for larger saccades.

**Figure 4.**
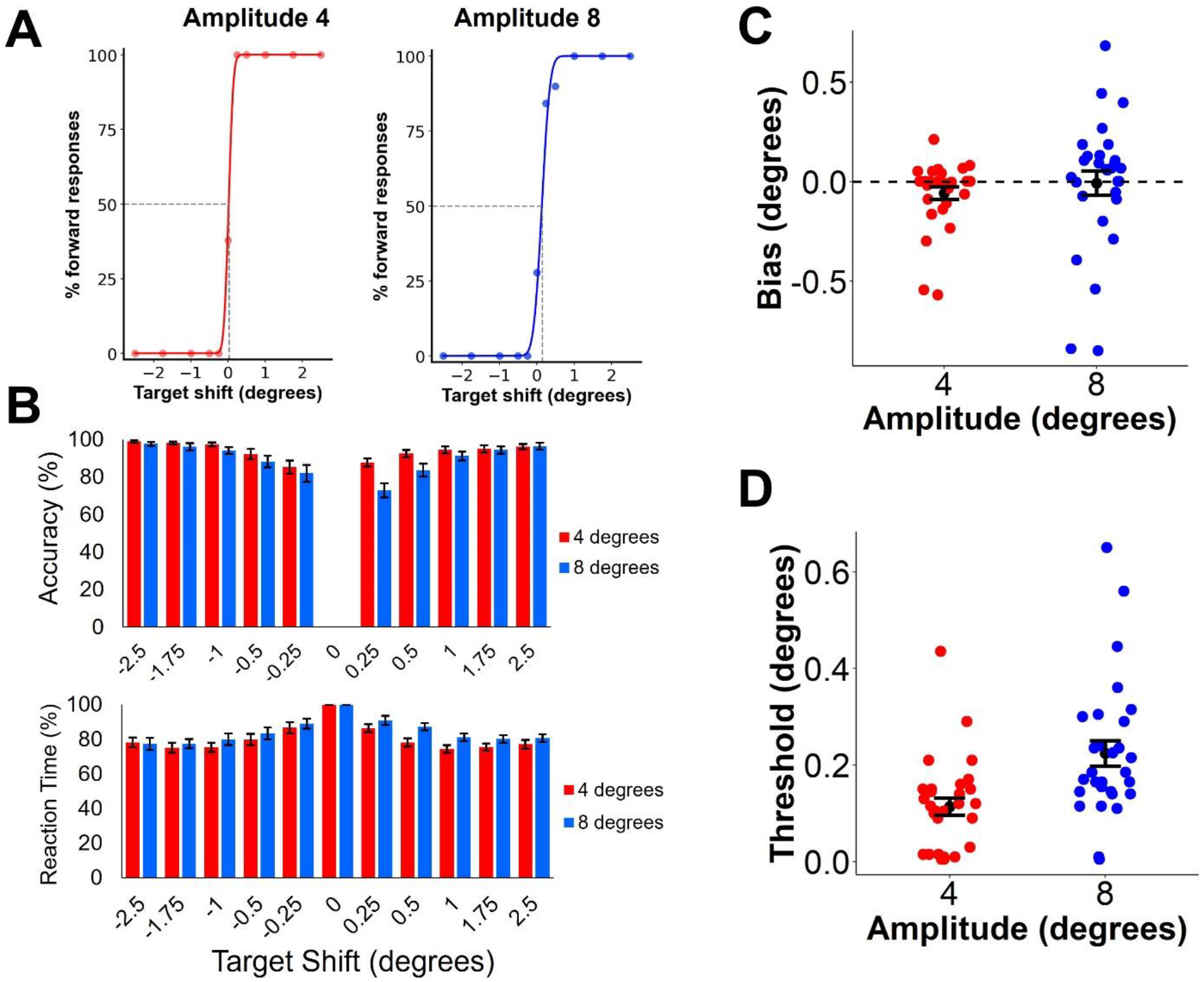
Behavioral and Psychometric Results Across Amplitude 4° and 8° Trials. (A) Psychometric curves (continuous line) fitted to the percentage of forward response data (individual filled circles) from a single subject across 4° (in red) and 8° trials (in blue). The bias is indicated with dashed lines and corresponds to the point on the psychometric curve with 50% forward responses. (B) Accuracy in participants’ manual decision of the shift direction (encoded as a percentage out of the total number of trials) and manual reaction time relative to each participant’s reaction time at the 0° target shift across target shifts for the 4° (in red) and 8° trials (in blue). (C) Psychometric bias spread across the 4° (in red) and 8° trials (in blue). Individual subjects are indicated with colored filled circles, black filled circles represent the mean across subjects, and the black error bars represent the standard error of the means. (D) Psychometric threshold across the 4° (in red) and 8° trials (in blue). Individual subjects are indicated with colored circles, black filled circles represent the mean across subjects, and the black error bars represent the standard error of the means.

A two-way analysis of variance (ANOVA) with repeated-measures and Greenhouse Geisser correction was performed to evaluate the effects of saccade amplitude and target shift on response accuracy. ANOVA results pertaining to response accuracy are reported in Table 3^c^^,d,e^. The main effect of saccade amplitude^c^ was due to a greater proportion of accurate responses following small saccade jumps (4° *M* = 94.33 %, *SE* = 2.04) relative to large saccade jumps (8° *M* = 90.58 %, *SE*= 3.00). There was a main effect of target shift mainly driven by progressively smaller proportions of correct responses for short target shifts relative to long target shifts (*p* < 0.001^d^). In addition, we found a significant interaction across the saccade amplitude and target shift factors^e^. Post-hoc pairwise *t*-tests run with a Bonferroni correction revealed key saccade amplitude effects at small forward shifts such as +0.25° (4° *M* = 87.70 %, *SE* = 2.02; 8° *M* = 72.83 %, *SE* = 3.77), *t* (28) = 4.80, *p* < 0.001^f^, but no difference for large shifts (e.g., -2.50°^g^ and +2.50°^h^, *p* = 1.0^g,h^).

**Table 3.** Behavioral Summary of ANOVA Results for Accuracy and Reaction Time Across Saccade Amplitudes and Target Shifts.

| Factor | *df* | *F* | *p* | $\eta_p^2$ |
| --- | --- | --- | --- | --- |
| **Accuracy (%)** |  |  |  |  |
| Saccade Amplitude | 1, 28 | 34.01 | < 0.001<sup>c</sup> | 0.55 |
| Target Shift | 1.86, 52.03 | 10.74 | < 0.001<sup>d</sup> | 0.28 |
| Interaction | 3.36, 94.04 | 6.13 | < 0.001<sup>e</sup> | 0.18 |
| **Reaction Time (%)** |  |  |  |  |
| Saccade Amplitude | 1, 28 | 8.14 | 0.008<sup>i</sup> | 0.23 |
| Target Shift | 2.84, 79.56 | 22.99 | < 0.001<sup>j</sup> | 0.45 |
| Interaction | 5.64, 157.79 | 2.92 | 0.12<sup>k</sup> | 0.09 |

A two-way analysis of variance (ANOVA) with repeated-measures and Greenhouse Geisser correction was performed to evaluate the effects of saccade amplitude and target shift on reaction time. ANOVA results pertaining to reaction time are reported in Table 3^i^^,j,k^. The ANOVA yielded a statistically significant main effect of saccade amplitude^i^ and was due to shorter reaction times following small saccade jumps (4° *M* = 80.59 %, *SE* = 2.87) relative to large saccade jumps (8° *M*= 84.31 %, *SE* = 2.90). There was a main effect of target shift^j^ mainly driven by progressively longer reaction times for short target shifts (e.g., -0.25°) relative to long target shifts (*p* < 0.001^j^). Finally, there was a significant interaction^k^ across the saccade amplitude and target shift factors. Post-hoc pairwise *t*-tests run with a Bonferroni correction revealed key saccade amplitude effects at small forward shifts such as +0.50°, *t* (28) = -4.44, *p* < 0.001^l^, but no difference for large shifts (e.g., -2.50°^m^ and +2.50°^n^, *p* = 1.0^m,n^).

### Saccadic Eye Movement Measures

We were interested in the influence of the visual error (VE) participants experienced on each trial. Fig. 5A provides an illustration of an example participant’s eye fixations following the initial saccade for the four possible target locations. There was a qualitatively larger spread for larger eye movements. We assessed potential visual gain differences across saccade amplitudes with a two-way repeated-measures ANOVA with Greenhouse Geisser correction^o,p,q^ (see Extended Data Figure 5-1A). The ANOVA (Table 4) yielded a statistically significant main effect of gain type^p^ with a greater proportion of undershoots (*M* = 70.01 %, *SE* = 3.42) relative to overshoots (*M* = 29.99 %, *SE* = 3.42). There was also a significant interaction^q^ which indicated a greater proportion of undershoots for saccades of amplitude 8° (*M* = 76.04 %, *SE* =3.34) relative to 4° (*M* = 63.98 %, *SE* = 3.20; see Extended Data Figure 5-1A).

**Figure 5.**
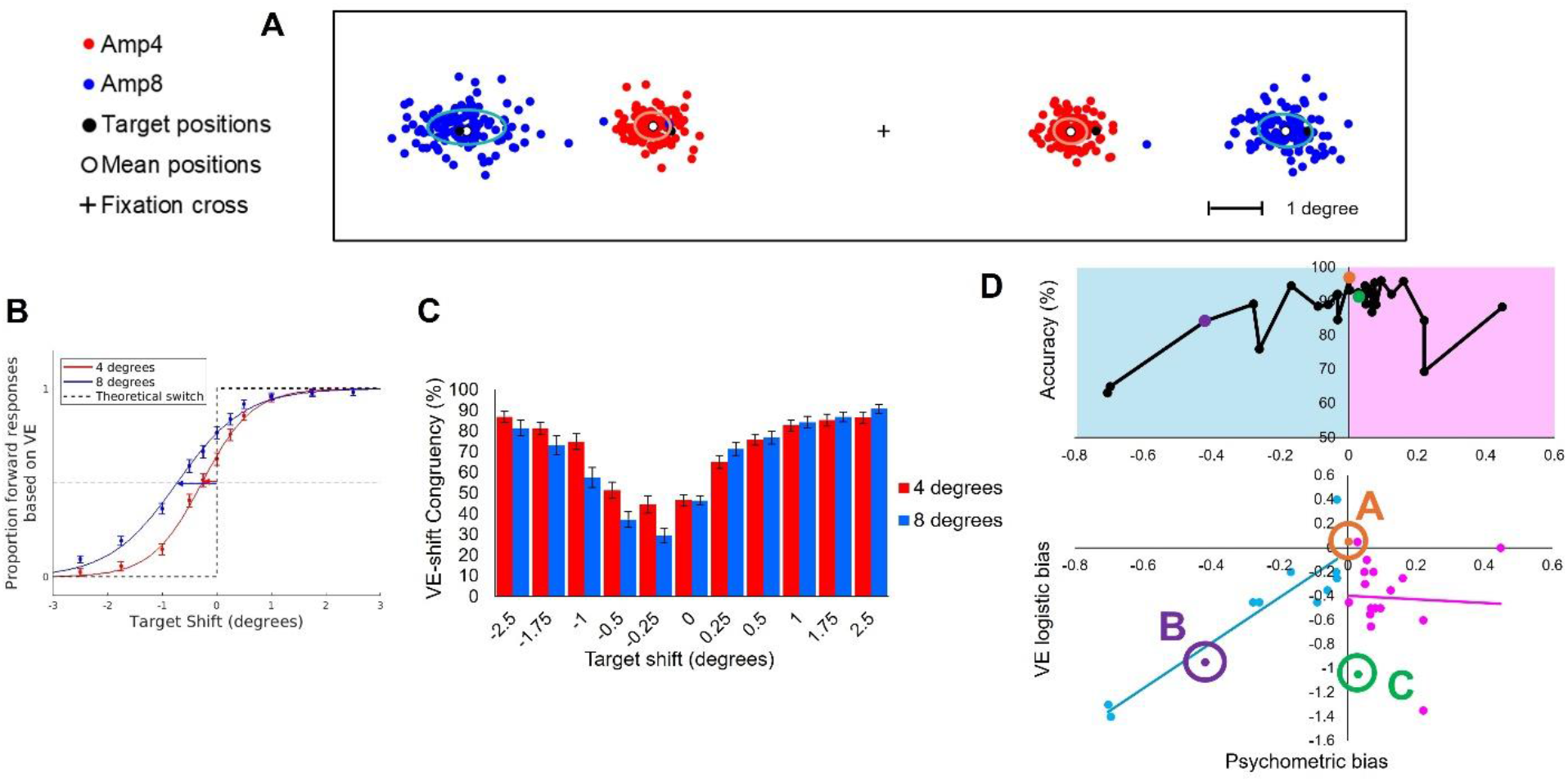
Illustration of Eye Fixations, VE-predicted Forward Responses, Bias Comparison, and Shift-VE Congruency Across Trials with Saccade Jumps of 4° and 8°. (A) Example of eye fixations after the initial saccade across trials in an individual participant. Red filled circles correspond to individual trials requiring a 4° saccade to the left or right of the fixation cross. Blue filled circles correspond to individual trials requiring a 8° saccade to the left or right of the fixation cross. Black filled circles indicate the target positions (left/right and 4°/8°). White circles indicate the mean positions within the corresponding eye fixation clusters. Colored ellipses indicate the spread around the mean positions in terms of 1 standard deviation of a normal Gaussian. (B) Examples of group-level VE-predicted forward responses across the backward and forward direction. A logistic function shows the best fit relative to VE-predicted forward responses across trials involving saccade amplitudes of 4° (in red) and 8° (in blue). A step function shown with a dashed line corresponds to what would have been expected had the eye saccade involved no bias. (C) VE-shift sign congruency across trials involving eye saccades of amplitude 4° (in red) and 8° (in blue). (D) Correspondence between the psychometric bias derived from button press responses and the VE-based logistic bias. Two groups of subjects have been identified: Turquoise filled circles indicate a linear relationship between the two biases along with the line of best fit. Pink filled circles show no linear relationship between the two biases as is indicated by the larger spread around the line of best fit. Three participants A, B, and C have been identified corresponding to three cases of no bias, a correspondence between the psychometric and VE bias, and a VE-only bias respectively. The inset plot shows individuals’ correspondence between their accuracy scores and psychometric bias ranked from most negative bias to most positive bias.

**Table 4.** Perceptual Summary of ANOVA Results for Gain Across Saccade Amplitudes and Gain Types and for VE-Shift Congruency Across Saccade Amplitudes and Target Shifts.

| Factor | *df* | *F* | *p* | $\eta_p^2$ |
| --- | --- | --- | --- | --- |
| **Gain** |  |  |  |  |
| Saccade Amplitude | 1, 28 | 0.00 | 0.99<sup>o</sup> | < 0.01 |
| Gain Type | 1, 28 | 38.76 | < 0.001<sup>p</sup> | 0.58 |
| Interaction | 1, 28 | 50.34 | < 0.001<sup>q</sup> | 0.64 |
| **VE-shift congruency** |  |  |  |  |
| Saccade Amplitude | 1, 28 | 14.26 | < 0.001<sup>s</sup> | 0.34 |
| Target Shift | 2.66, 74.50 | 83.28 | < 0.001<sup>t</sup> | 0.75 |
| Interaction | 5.91, 165.47 | 13.13 | < 0.001<sup>u</sup> | 0.32 |

A one-way repeated-measures ANOVA with Greenhouse Geisser correction yielded a main effect of saccade amplitude on VE bias, *F* (1, 28) = 40.31, *p* < 0.001^r^, *η^2^* = .21, with a larger, more negative VE bias at saccadic amplitude 8° (*M* = - 0.75°, *SE* = 0.12°) compared to 4° (*M* = - 0.24°, *SE* = 0.05°). This negative VE biasing effect is believed to mostly be the result of movement undershoots (see Fig. 5B and Extended Data Figure 5-1A), which may significantly contribute to incorrect VE-based responses (see Fig. 2A).

We next assessed potential VE and target shift congruency differences across target shifts and saccade amplitudes. A two-way repeated-measures ANOVA with Greenhouse Geisser correction (Table 4) yielded a statistically significant main effect of saccade amplitude^s^ with more congruency across VE and target shift for 4° (*M* = 71.00 %, *SE* = 4.20) relative to 8° saccades (*M* = 66.83 %, *SE* = 4.92). The ANOVA also revealed a main effect of target shift^t^ which was driven by more incongruent VEs for short backward target shifts (e.g., -0.25°) relative to large target shifts (*p* < 0.001^t^). Finally, a significant interaction revealed that trials involving small backwards target shifts yielded lower levels of VE-shift congruency following 8° saccades relative to 4°, but there were no VE-shift congruency differences across saccade amplitudes for large forward shifts (*p* = 1.0^u^; see Fig. 5C).

One key comparison was the relationship between the empirical psychometric bias based on manual responses and VE-predicted forward responses (see Fig. 5B). The comparison and corresponding correlations are shown in Fig. 5D. The main finding was that the two biases were most correlated in the negative (backward) direction, *r* (27) = 0.92, *p* < 0.001^v^, which fits with the generally negative VE bias across individuals (see Fig. 5B) and with participants’ propensity to undershoot the visual target (see Figure Extended Data Figure 5-1A). On the other hand, there was no significant correlation between the two biases in the positive (forward) direction, *r* (27) = - 0.28, *p* = 0.14^w^ (see Fig. 5D). When considering accuracy scores across the groups with a positive and negative psychometric bias, we found a statistically significant difference between the two groups, *t* (28) = 2.24, *p* = .03^x^, with individuals with a positive psychometric bias generally demonstrating a greater accuracy (*M* = 90.56%, *SE* = 1.49) relative to individuals with a negative psychometric bias (*M* = 83.45%, *SE* = 3.25). When plotting accuracy against psychometric bias, we found that individuals with a progressively less negative psychometric bias also had increasingly higher accuracy scores (see Fig. 5D and Extended Data Figure 5-1B). On the other hand, individuals with a positive psychometric bias tended to have generally higher accuracy scores regardless of the magnitude of their bias (see Fig. 5D and Extended Data Figure 5-1B).

### VE coefficient and Characterization of the Psychometric Bias

Fig. 6 shows the relationship between the VE bias and the resulting biasing effects on perceptual judgments. Note that the VE coefficients indicate at which target shifts an incongruency in VE relative to the direction of the target shift may lead to incorrect responses. Participant A corresponds to an example subject with close to no bias (along the psychometric and VE dimensions; see Fig. 5D) and a well-centered VE coefficient centroid. Participant B and Participant C both illustrate example subjects with a significant negative (backwards) VE bias. However, only participant B appeared to make incorrect responses, thus resulting in a more negative centroid. Conversely, participant C did not make as many errors resulting in low VE coefficients and a more centered VE coeff centroid. Overall, the VE coeff centroid was found to be strongly correlated with the psychometric bias, *r* (27) = 0.90, *p* < 0.001^y^ (see Fig. 8A). Further assessment of the statistical robustness of the VE coeff centroid against baseline noise uncovered an SNR = 3.27 (*SE* = 0.79) suggesting that the centroid is a reliable estimate (SNR > 3.00).

**Figure 6.**
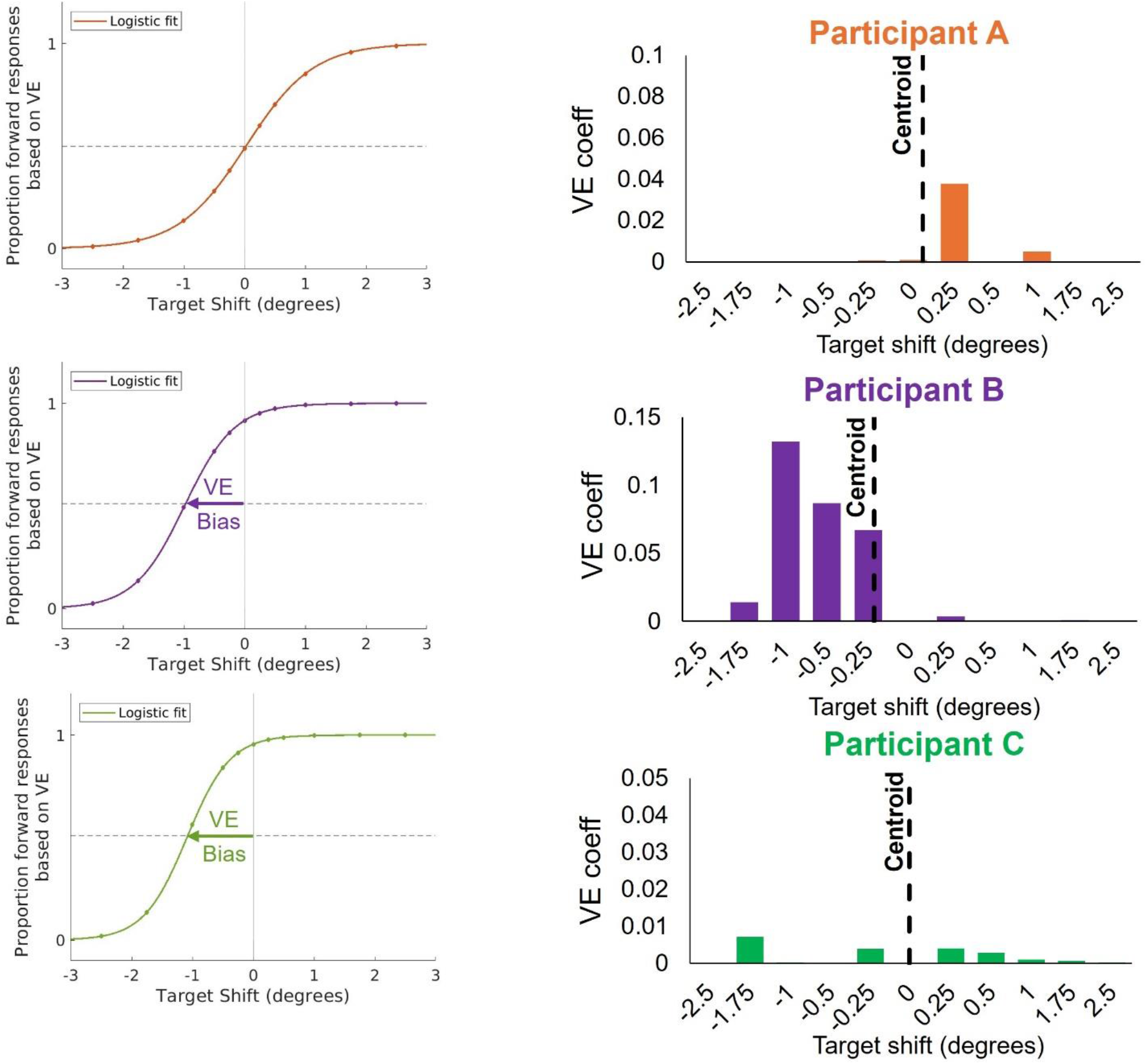
Biasing Profiles and VE Contribution to Bias Across Three Participants. (A) VE-predicted forward responses in the same three characteristic participants A (top), B (middle), and C (bottom). A logistic function fit to the VE-predicted forward responses can be used to identify the global VE bias. Participant A’s fit is shown in orange, participant B’s fit is shown in purple, and participant C’s fit is shown in green. (B) VE coefficient distribution across the same participants A (top), B (middle), and C (bottom). The VE coefficient centroid has been identified with a dashed line. (C) Quantification of the VE bias in the three participants A, B, and C. (D) Quantification of the psychometric (VE+CD) bias in the three participants A, B, and C.

To assess the relationship between the RT deviation from symmetry and the VE coefficients, we employed a linear mixed-effects model (LMEM). The analysis revealed a significant effect of the VE coefficient on RT (VE_coeff *γ* = 366.44*, p* < 0.001^z^; CI = [267.63, 463.68]). These results demonstrate that the VE coefficients are suitable to mechanistically account for decision time biases as part of the DDM framework.

### DDM Individual Fits of Choice Data and Reaction Time

The next step was to use the VE coefficients to simulate the effects of a participant’s bias on their decision-making process in the visual perception task. The VE was used as an input in the DDM along with choice (% forward responses) and manual RT data, and directly led to a modulation of the drift rate as illustrated on Fig. 3B. For each participant and each saccade amplitude, the model optimized the 4 DDM parameters: *r, DB*, *B*, and *NDT*. In addition, a final weight was estimated to determine the extent of drift rate bias at the level of the inputs (see equation 7). Two separate parameter sets were estimated for each saccade amplitude. Fig. 7 provides detailed results for the three representative participants A, B, and C, and presents their corresponding psychometric and RT profiles along with their DDM model fits. As a reference, we also show a uniform version of the DDM that assumed symmetry across forward and backward target shifts.

**Figure 7.**
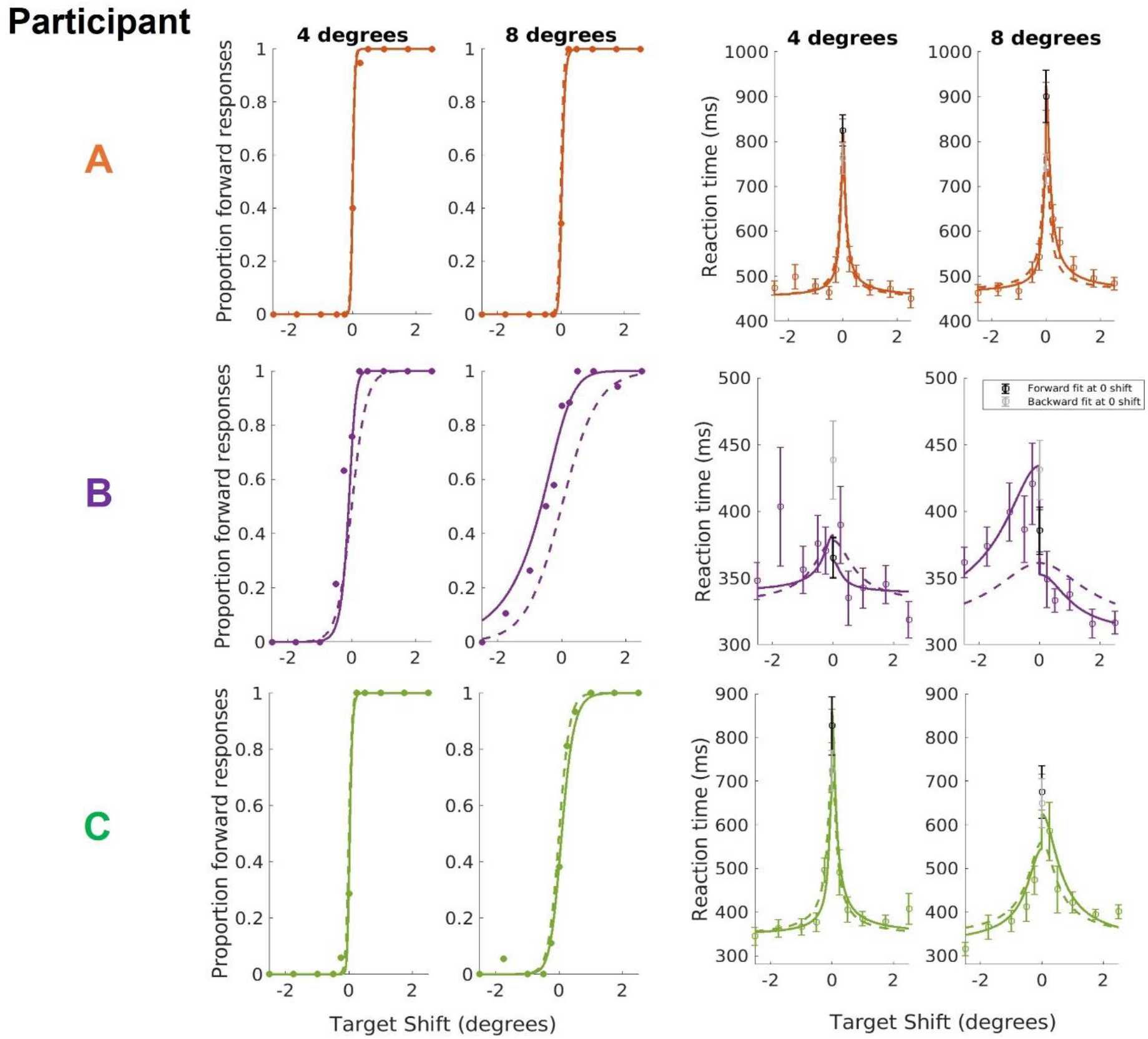
Psychometric and Reaction Time Curve Fits Based on DDM Simulations and Relationship between VE Coeff Centroid and Degree of RT asymmetry. Three participants’ psychometric and reaction time curves were fitted to the data split by target shift based on DDM simulations across 4° and 8° trials. Manual reaction time data differentiate between forward (in black) and backward reaction times (in gray) at the 0° target shift. Dashed lines correspond to DDM simulations with the uniform baseline model. Solid lines correspond to DDM simulations from the model that incorporates a bias term and a VE-based weight. For reaction time data, circles indicate the mean, and the error bars indicate the standard error of the means. Participant A corresponds to an example of a VE coeff centroid being close to 0° and no VE bias, and is shown in orange at the top. Participant B corresponds to an example of a negative VE coeff centroid and a negative VE bias, and is shown in purple in the middle. Participant C corresponds to an example of a VE coeff centroid being close to 0° and a negative VE bias, and is shown in green at the bottom. On the right, individuals’ VE coeff centroid are plotted against their degree of RT asymmetry. Quantification of both the VE coeff centroid and the degree of RT asymmetry across participants A, B, and C is illustrated in the middle and bottom.

Specifically, participant A had a VE coefficient centroid close to 0°. Their psychometric curves and RT profiles displayed symmetry across the forward and backward shift trials. Furthermore, this symmetric relationship was also reflected by the significant overlap between the psychometric and reaction time profiles across the two types of DDM simulations. Participant A’s low psychometric bias was also reflected in a low RT asymmetry χ^2^ (see Fig. 7).

Participant B was characterized by a negative VE coefficient centroid. Their psychometric curves displayed greater proportions of forward responses for backward target shifts indicating incorrect decisions along the backward direction. The negative VE coefficient centroid was also significantly related to a greater RT asymmetry χ^2^ (see Fig. 7). In terms of manual RT, we found that the participant’s data fit better with the VE-driven DDM, with slower reaction times along the backward direction relative to forward target shifts. Participant B’s DDM-based VE coefficient was estimated as a scalar of large magnitude and was consistently negative across saccade amplitude, reflecting the negative VE bias contribution to decision-making.

Lastly, participant C was characterized by a negative (backward) VE bias but insignificant VE coefficient centroid and low RT asymmetry χ^2^ (see Fig. 7). Yet, their psychometric curve displayed the same symmetry that was characteristic of participant A with no bias, and their manual RT profiles also displayed the same symmetry as participant A. Overall, this means that even though participant C’s VE profiles would have led to a predicted perceptual bias, this was not found to be the case at the level of the actual psychometric curves. This was also reflected in the distribution of VE coefficients (see bottom row of Fig. 6).

### Inter-individual Differences in Centroid & RT Asymmetry and Correlations

Across all subjects, the RT asymmetry χ^2^ was inversely related to the VE coefficient centroid and psychometric bias, *r* (27) = - 0.69, *p* < 0.001^aa^ (see Fig. 8B). This result indicates that the participants that tended to have a greater RT asymmetry also tended to have a more biased VE coefficient centroid. This also suggests that greater asymmetry in the psychometric and RT profiles are strongly correlated to larger contributions of the VE coefficients towards the drift rate offset (i.e., an influence on the perceptual response, see equation 7).

**Figure 8.**
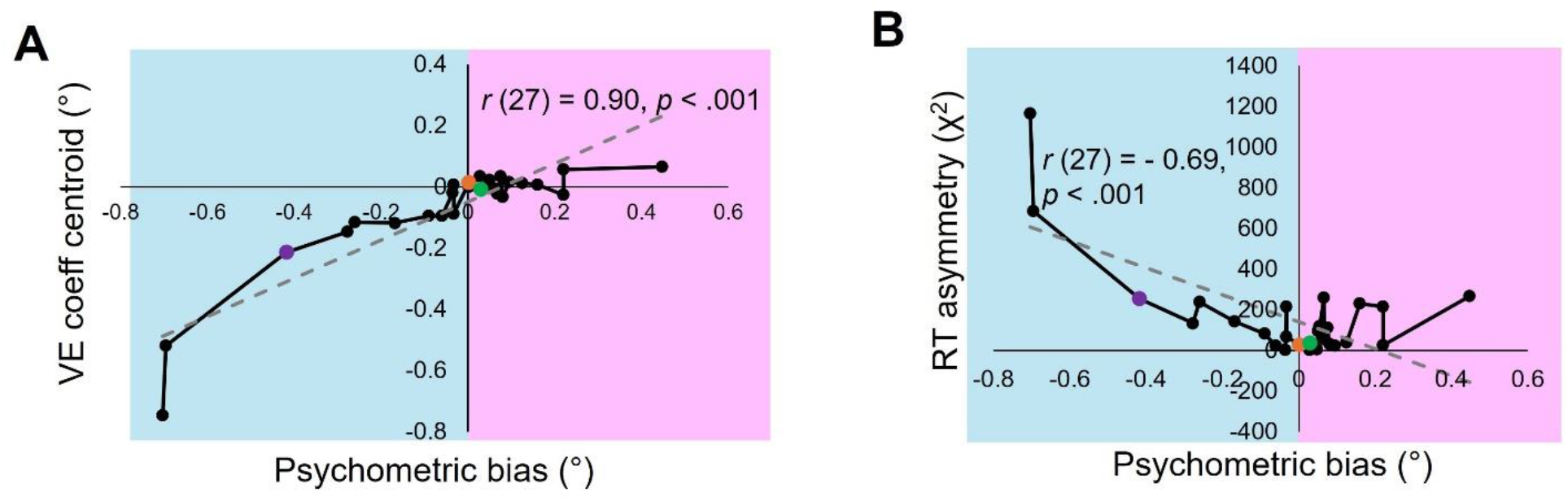
Interindividual relationships in VE coeff centroid and RT asymmetry. (A) Individuals’ correspondence between their VE coefficient centroid and psychometric bias ranked from most negative bias to most positive bias. The blue panel highlights individuals with a negative psychometric bias and the pink panel highlights individuals with a positive psychometric bias. A linear relationship between both was identified with a dashed gray line of best fit. (B) Individuals’ correspondence between their RT asymmetry (χ^2^) and psychometric bias ranked from most negative bias to most positive bias. The blue panel highlights individuals with a negative psychometric bias and the pink panel highlights individuals with a positive psychometric bias. Three participants A, B, and C have been identified corresponding to three cases of no bias (orange), a correspondence between the psychometric and VE bias (purple), and a VE-only bias (green) respectively. A linear relationship between both was identified with a dashed gray line of best fit.

When considering VE coefficient centroid across the two groups with a positive and a negative psychometric bias, we found a statistically more negative centroid in the group with a negative psychometric bias (*M* = -0.20°, *SE* = .07), *t* (28) = 3.77, *p* < 0.001^bb^, relative to the group with a positive psychometric bias (*M* = 0.01°, *SE* = 0.01; see Extended Data Figure 5-1C and Fig. 8A). When plotting the centroid against participants’ psychometric rank, we found that individuals with a progressively less negative psychometric bias tended to have a generally less negative centroid. On the other hand, individuals with a positive psychometric bias were found to generally have a VE coefficient centroid around 0 (see Fig. 8A).

In terms of the inter-individual differences in RT asymmetry, we found a statistically more significant χ^2^ statistic in the group with a negative psychometric bias (*M* = 272, *SE* = 105), *t* (28) = 2.11, *p* = .04 ^cc^, relative to the group with a positive psychometric bias (*M* = 91.61, *SE* = 21; see Extended Data Figure 5-1D and Fig. 8B). Across subjects, the RT distribution transitioned from generally asymmetric (i.e., high χ^2^ statistic) to symmetric (i.e., low χ^2^ statistic) as the psychometric bias went from negative to positive (see Fig. 8B).

## Discussion

In this study, we leveraged the DDM framework to characterize the evidence accumulation process that underlies the perception of visual stability across saccadic eye movements. One key new feature was the incorporation of the visual error (VE) into the model to account for the visuospatial biasing effects of sensory information on decision making. We adapted a trans-saccadic shift detection task (Bansal et al., 2018; Bansal and Joiner, 2021) and examined participants’ decision-making behavior. We measured perceptual sensitivity in terms of the psychometric bias and the psychometric threshold, and we operationalized decision-making behavior in terms of reaction time (RT) and response accuracy. We expected small target shifts to induce a decision-making bias as would be predicted from the saccadic suppression of displacement (SSD). Computationally, we focused on the hypothesis that by incorporating the VE the DDM would capture idiosyncratic biases in perception and RT profiles.

### Expansion of Perceptual Effects of SSD to Decision-Making

Psychometric analyses replicated previous findings on trans-saccadic perception (Joiner et al., 2013; Bansal et al., 2015; Jayet Bray et al., 2016; Bansal and Joiner, 2021). Specifically, the psychometric threshold proportionally increased with the initial saccade’s amplitude and smaller target shifts were associated with lower accuracy and slower RT (Fig. 4). Both results provide further evidence that small target shifts required longer decision times relative to larger target shifts and were more difficult to detect after longer saccades. These results fit with the main assumptions of the DDM, which posits that an increase in the target shift magnitude should be associated with a monotonic increase in accuracy and decrease in RT (Smucny et al., 2023b).

Relevant for our subsequent computational modeling simulations, we also observed a well-known undershooting tendency which led participants to fall short of the intended visual targets (Fig. 5 and Extended Data Figure 5-1A). This occurred to a greater extent following larger saccadic eye movements which aligns with previous experimental reports (Fig. 5; Bansal et al., 2018; Collewijn et al., 1988; Leigh and Zee, 2015). Because of the SSD, it could be the case that small backward target shifts directly following the saccade also impacted perceptual sensitivity, particularly around the eye endpoint location. (Fig. 4 and Fig. 5C). In such cases, participants must rely on extraretinal information in addition to the VE to determine the correct direction of the target shift. This mechanism has previously been described in terms of Bayesian inference mechanisms, specifically in the prefrontal cortex (Lange et al., 2021; Goris et al., 2024; Langlois et al., 2025)

In the trans-saccadic task, a greater weight on the VE would result in incorrect perceptual judgments most likely due to the down-weighing of extraretinal signals (see Fig. 2A). We found that participants with a negative psychometric bias tended to make perceptual judgments that were more directly correlated with predictions from the VE. In addition, perceptual accuracy tended to systematically increase as individuals’ psychometric bias became less negative. Conversely, individuals with positive or no psychometric bias deviated strongly from the direct predictions based on the VE, suggesting the incorporation of extraretinal signals (Fig. 5). Except for occasional outliers demonstrating a particularly low VE bias and low perceptual accuracy, we found that individuals with a positive psychometric bias tended to display higher perceptual accuracy levels relative to individuals with a negative psychometric bias regardless of magnitude.

We computed coefficients related to the VEs that estimated participants’ propensity towards VE-driven incorrect perceptual judgments (Fig. 2). Within participants, such VE coefficients could be characterized in the form of a distribution whose center of gravity (the centroid) provided a direct quantification for individuals’ inclination towards such VE-driven incorrect perceptual errors. At the group level, this metric was strongly related to both the psychometric bias (Fig. 6) and RT asymmetry (Fig. 7), and distinguished individuals with similar VE profiles (i.e., participant B and C). We also found that the VE coefficients could successfully predict individuals’ manual RT, thus capturing decision-making temporal biases during the course of evidence accumulation.

### DDM Framework & Perceptual Bias on Decision-Making

We found that the DDM was able to account for patterns in asymmetry as described above, both in terms of manual choice and RT. In terms of model design, the key novelty was the addition of sensory information to the model. To simulate patterns of asymmetry, key design choices were the offset of the decision process’s starting point with a bias term and the modulation of target shifts’ drift rates, proportionally to the VE coefficients (Fig. 3). To assess fit with data, we compared the VE-driven DDM against a uniform version which was fully symmetric. In the latter, the decision time for backward target shifts matched the decision time for trials with forward target shifts of the same magnitude. When incorporating the VE-based coefficients into the DDM, the framework captured individual biases in manual responses and RT profiles (Fig. 7). Relative to single psychometric bias estimates, the DDM provided a more granular representation of individuals’ decision-making biases across target shifts by representing each shift’s corresponding drift rate offset. In addition, this modified model also included an explanatory mechanism that directly tied decision-making biases to perceptual properties related to the VE. That is, such biases were linked to cases in which there was (1) greater incongruency between the VE and the target shift direction and (2) a greater propensity towards incorrect perceptual judgments. Combined, this resulted in a meaningful alteration of the evidence accumulation process and subsequently the asymmetric psychometric functions and RT profiles.

In terms of inter-individual differences, we found that individuals with a negative psychometric bias were characterized by generally more negative VE coefficient distributions and by greater RT asymmetry relative to individuals with a positive psychometric bias. Importantly, both measures (the VE coefficient distribution and RT asymmetry) were significantly related to each other and also to the psychometric bias (Fig. 8). Based on individuals’ biases, we determined that the greatest patterns of behavioral asymmetry (i.e., perceptual judgment and reaction time profiles) systematically coincided with markedly negative psychometric biases (Fig. 8). This result further highlights the potential future use of the VE-based computational framework as a tool to identify meaningful inter-group differences, potentially across clinical groups characterized by differential weighing of the VE.

### Weight of the VE, Link to Corollary Discharge, and Relevance For Clinical Applications

One extraretinal signal that may influence perceptual decision-making in the trans-saccadic visual perception task is the corollary discharge (CD) signal. CD specifically refers to the internal copy of a motor command which is created at the time of movement execution (Kawato, 1999; Wurtz and Sommer, 2004; Tseng et al., 2007). CD is used to prepare the cortex for sensory inputs that would result from self-generated movement, and to distinguish self-induced sensory information from externally-generated sensory signals (Wolpert et al., 1995, 1998; Kawato, 1999). In this task, CD signals provide a spatial reference frame for understanding where visual targets shifted from relative to the saccadic eye movement (Bridgeman et al., 1975; Hall and Colby, 2011; Cavanaugh et al., 2016). In the brain, CD signals are processed via the superior colliculus (SC) – mediodorsal thalamus (MD) – frontal eye field (FEF) neural pathway (Sommer and Wurtz, 2008). To make an accurate judgment as to where the visual target was shifted to in the task, such CD signals are integrated with cortical visual inputs originating from the ventral and dorsal visual streams through the Lateral Intraparietal (LIP) cortex at the level of the FEF (Sommer and Wurtz, 2008). In the case of an undershoot, sign incongruencies between backwards target shifts and the VE’s direction are interpreted differently depending on the integration of visual inputs and CD signals. With an over-reliance on inputs from the visual processing streams, an incorrect perceptual judgment will ensue, as accurate perceptual judgments require actively incorporating the CD inputs from the SC-MD-FEF neural pathway with visual inputs from cortex.

From a clinical standpoint, previous research has found that individuals with schizophrenia tend to make perceptual decisions that are aligned with the judgments that would have been predicted had they solely relied on VE information (Bansal et al., 2018), Additionally, these patients demonstrate symptom-related disturbances in their ability to remap visual targets following saccades (Rösler et al., 2015; Thakkar et al., 2015). On the other hand, healthy control participants make perceptual judgments that deviate from VE-predicted responses, as would have been expected following correct integration of the VE with CD signals (Rösler et al., 2015; Thakkar et al., 2015; Bansal et al., 2018). Overall, perceptual estimates based on the VE may provide DDM-based behavioral markers for the investigation of sensory-based decision-making biases at the neural level (Shadlen and Newsome, 1996, 2001; Roitman and Shadlen, 2002; Mulder et al., 2014; Brody and Hanks, 2016; Cerracchio et al., 2023; Forstmann and Turner, 2024). In this experimental context, the VE-driven DDM provides a quantification of how evidence accumulation may be directly altered by overly relying on external sensory signals thus resulting in perceptual judgment errors and manual response and RT asymmetric patterns (e.g., schizophrenia, Butler et al., 2001; Martínez et al., 2008; Adámek et al., 2022, 2024). Future research may leverage such methods to determine to what extent individuals with schizophrenia may be biased by the processing of sensory information due to a potential downweighing of CD signals, as suggested by patients’ sensory gating disruptions and difficulties in filtering out irrelevant sensory information from the environment (Ford and Mathalon, 2004; Beño-Ruiz-de-la-Sierra et al., 2024).

## Conclusion

In summary, the present study leveraged the experienced sensory information (captured with the VE coefficients) to determine an individuals’ propensity towards sensory-driven incorrect perceptual judgments. Overall, the VE-driven DDM framework provides mechanistic insight into the direct effects of sensory information weighing on patterns of behavioral biases in perceptual judgments and RT profiles. Future research will investigate the utility of this new form of the DDM in clinical contexts to shed light on the underlying neural mechanisms involved in perceptually-based computations and sensorimotor learning in psychopathology.

## Author Contributions

WMJ acquired funding, WMJ and TAL designed research and acted in a supervisory role, CLE performed research, PGG analyzed data, WZ and TDH assisted with analyses, PGG wrote the paper, WMJ, TAL, TDH, and CSC provided feedback and made paper edits.

## Acknowledgements

We would like to thank Grace Shen and Staci Board for help with data collection.

## MULTIMEDIA, TABLES, AND EXTENDED DATA

**Extended Data 1. DDM Code For Preprocessing and Model Simulations.** Folder contains the MATLAB code that was used to preprocess the data and run the different DDM simulations of evidence accumulation leading to perceptual judgments. One model is the baseline DDM with no bias. A second model leverages visual error information to simulate VE-based patterns of asymmetry in manual choice and reaction time across individuals. All code and data are available at https://doi.org/10.6084/m9.figshare.30090127

**Extended Data Figure 3-1.**
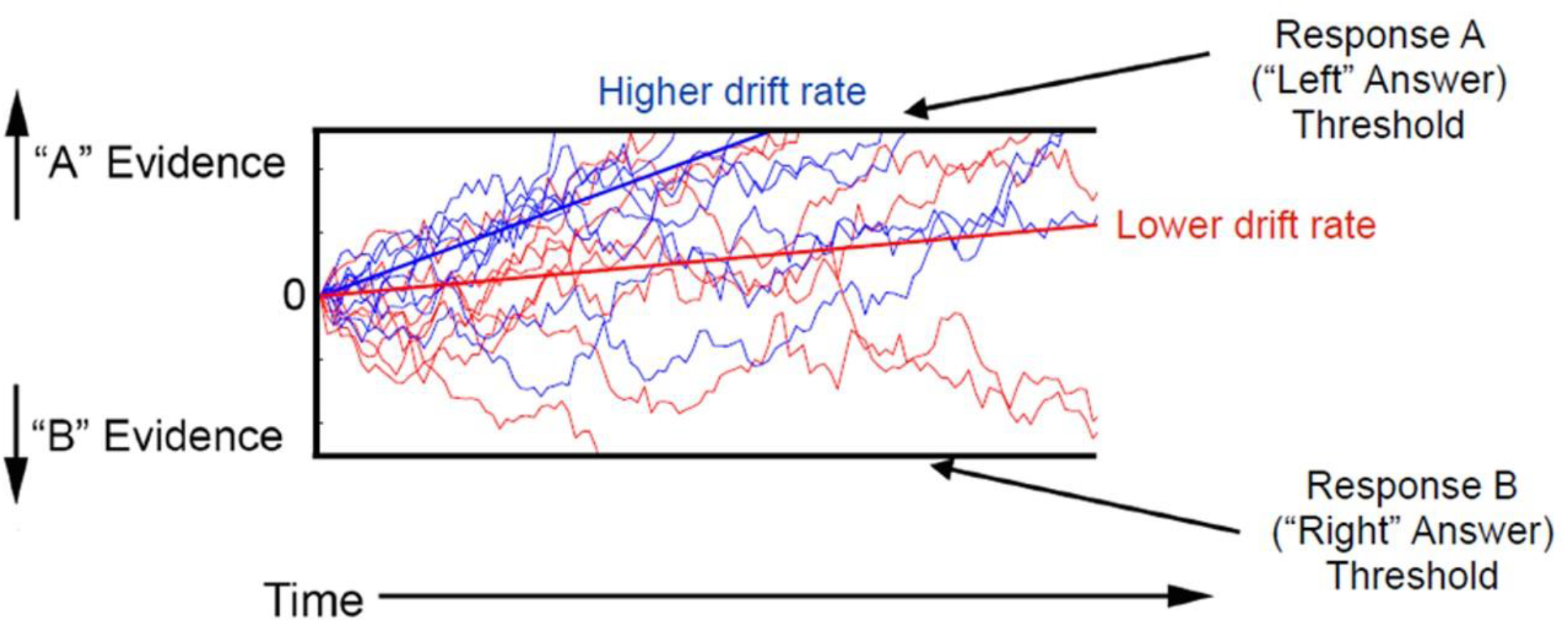
Visual Representation of the Drift Diffusion Model (DDM) and Drift Rate. The DDM represents the information accumulation process that unfolds during decision-making, and is modeled as a function of the signal-to-noise ratio of information gathering towards the decision point, a decision bound for making a decision, and non-decision time (due to motor processing). The drift rate is the average slope of evidence accumulation (represented with solid lines). On this diagram, “A” and “B” correspond to each of the two possible shift directions (‘left’ and ‘right’).

**Extended Data Figure 3-2.**
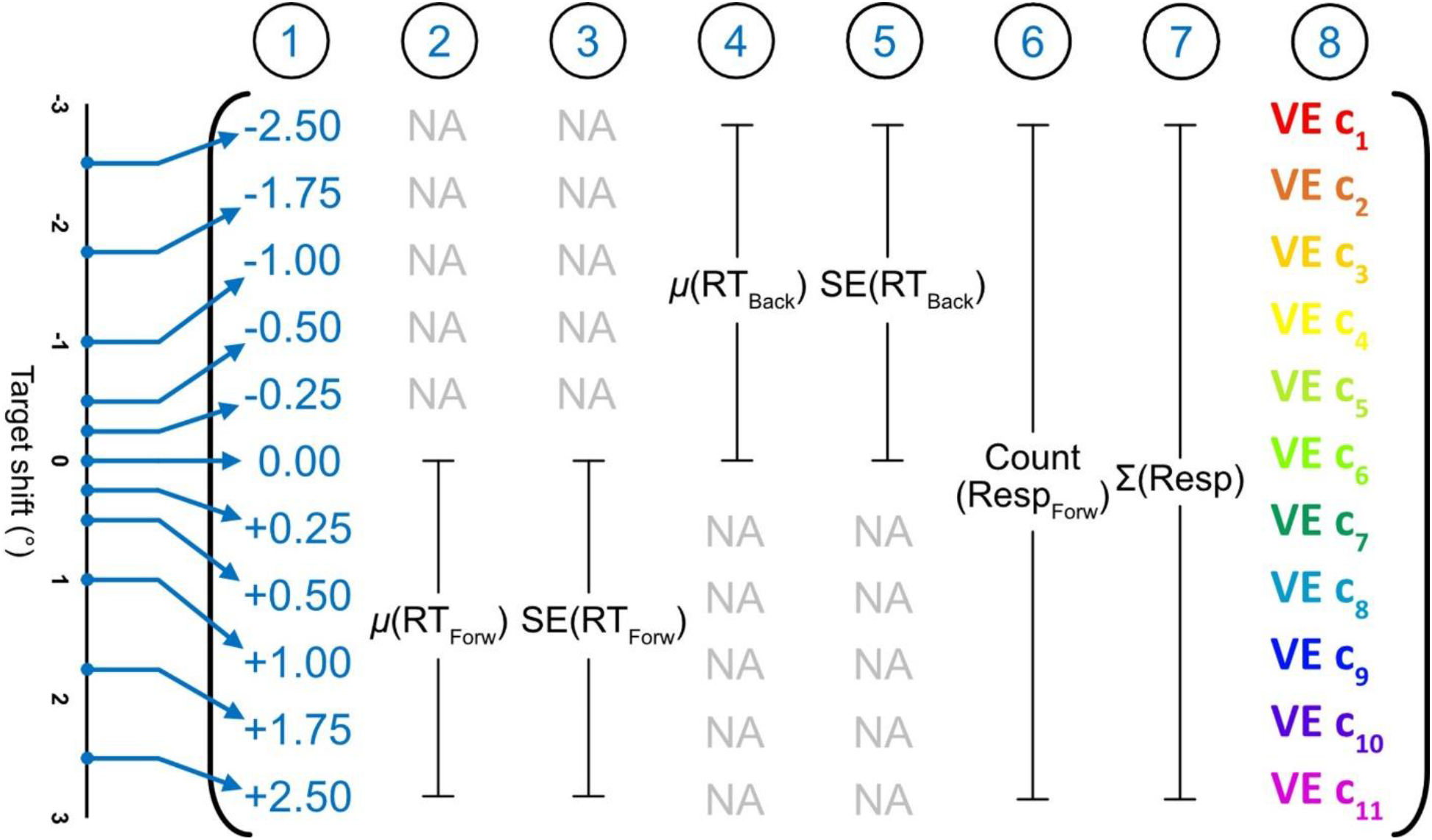
Visual Representation of the DDM Input Matrix. Data pertaining to the target shifts, reaction time, manual choices, and VE coefficients were entered into an input matrix, and were the basis of DDM analyses. In this matrix, each of the 11 rows corresponded to trials with different target shifts. There were a total of 8 columns. The first column indicated the target shift. The second and fourth columns indicated the mean manual reaction time across forward trials (2^nd^ column) and backward trials (4^th^ column). The third and fifth columns indicated the manual reaction time standard errors across forward (3^rd^ column) and backward trials (5^th^ trials). The sixth and seventh columns indicated the total number of forward responses and the total number of responses respectively. The eighth column indicated the VE coefficients that were estimated based on the proportion of incorrect responses and VE-predicted forward responses.

**Extended Data Figure 5-1.**
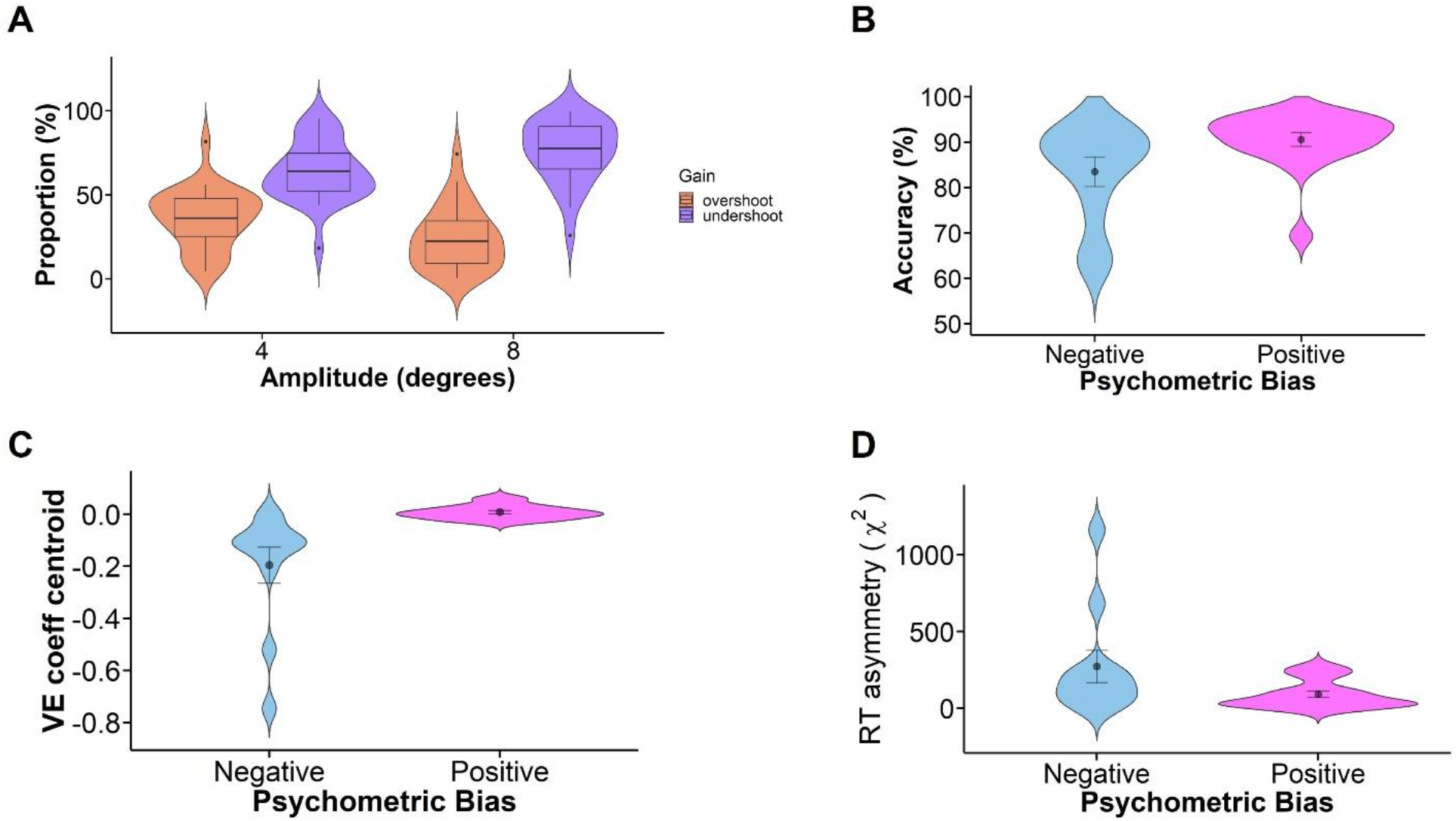
Illustration of the Gain Across Saccade Jumps of 4° and 8°. (A) Proportions of trials with overshoots and undershoots computed in terms of the gain relative to initial target position. Gain was calculated separately across 4° and 8° trials. Distributions of gains are shown with a violin plot and box plot. Brown distributions correspond to overshoots. Purple distributions correspond to undershoots. (B) Violin plots displaying the distribution of accuracy scores across negative and positive psychometric groups. Black filled circles indicate the group means, and the error bars correspond to standard errors of the mean. (C) Violin plots displaying the distribution of VE coefficient centroids across negative and positive psychometric groups. Black filled circles indicate the group means, and the error bars correspond to standard errors of the mean. (D) Violin plots displaying the distribution of RT asymmetry (χ^2^) across negative and positive psychometric groups. Black filled circles indicate the group means, and the error bars correspond to standard errors of the mean.

