## Supplementary figures and images for "Computational Characterization of Decision Making During Trans-saccadic Visual Perception"

### Extended Data Figure 3-1

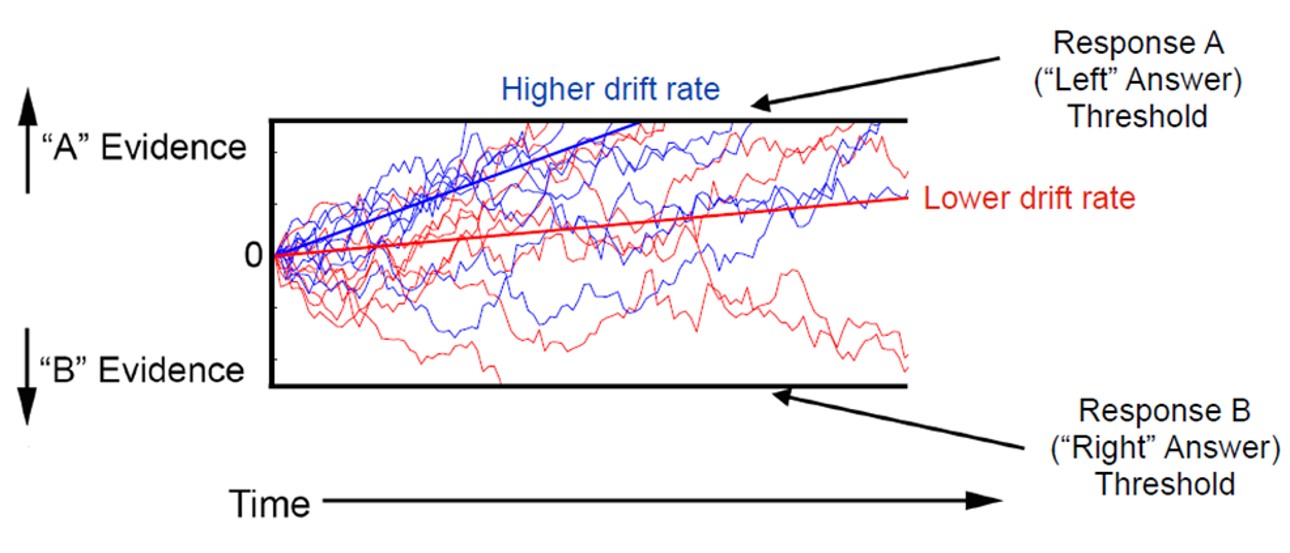

### Extended Data Figure 3-2

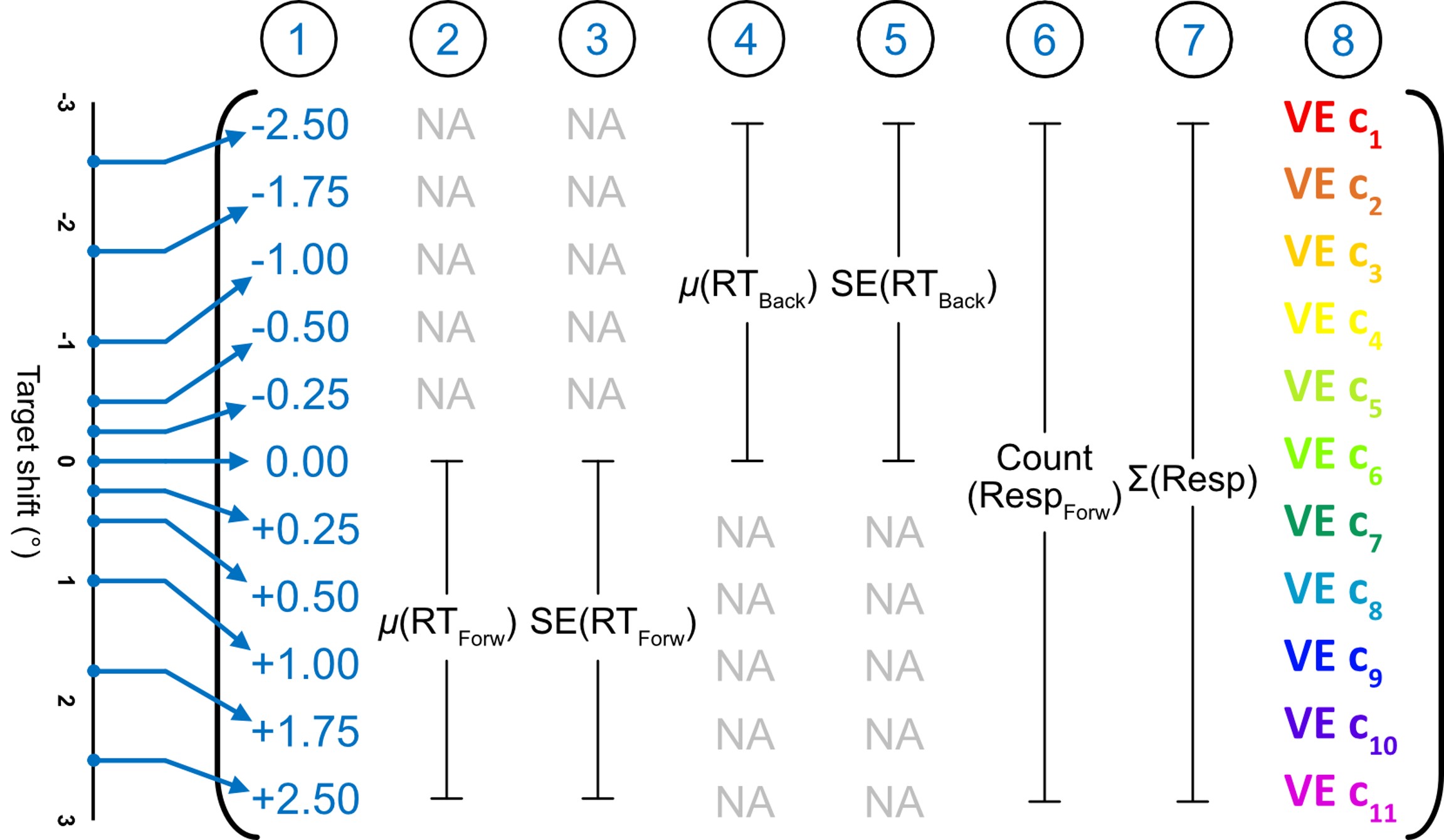

### Extended Data Figure 5-1

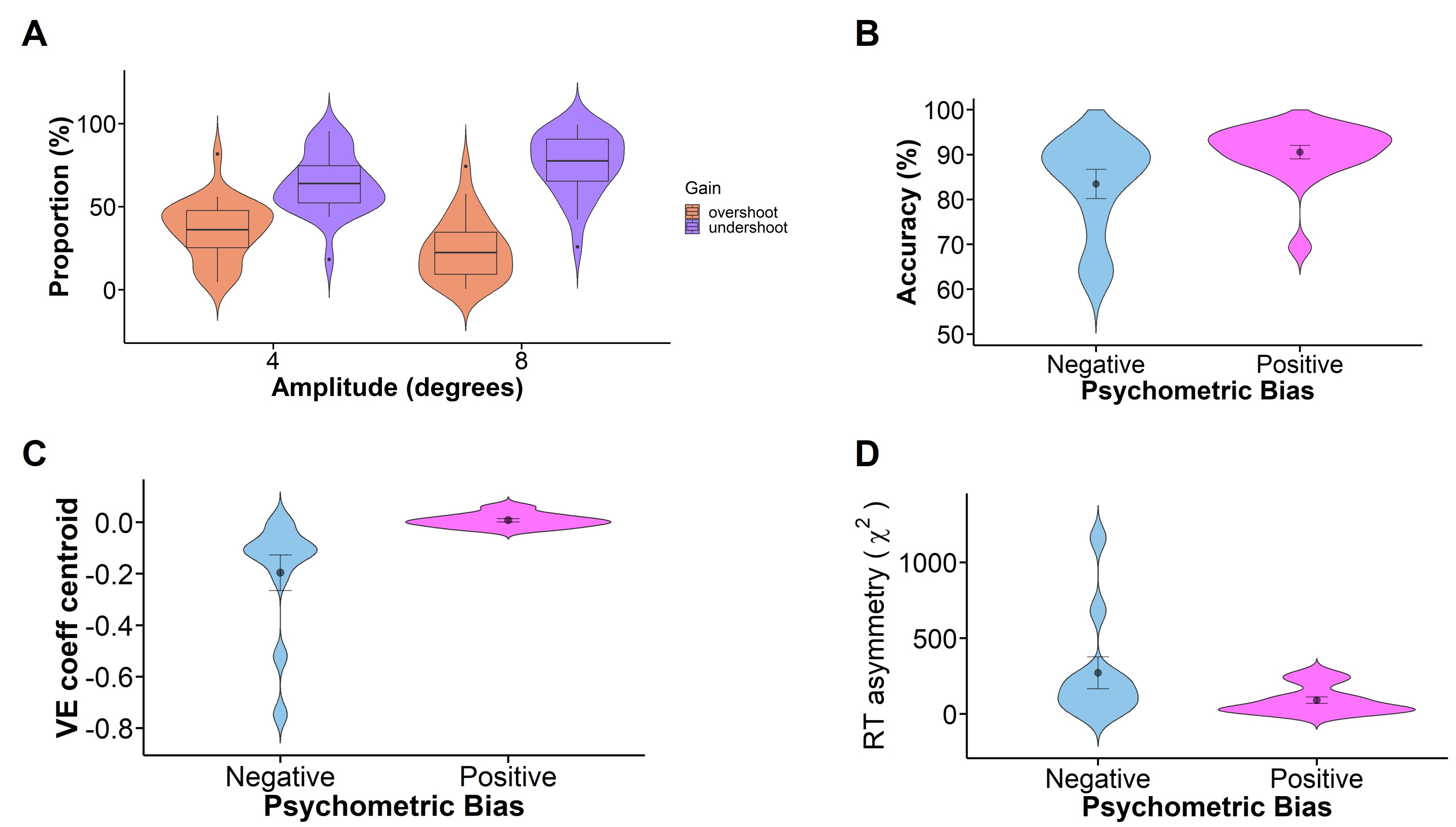
